# Entanglement dilution and high fractal dimension mediated by loop extrusion revealed in simulations of active polymer melts

**DOI:** 10.64898/2026.08.08.743709

**Authors:** Brian Chan, Michael Rubinstein

**Affiliations:** Thomas Lord Department of Mechanical Engineering and Materials Science, Duke University, Durham, North Carolina, 27708, United States; Departments of Biomedical Engineering, Physics, and Chemistry, Duke University, Durham, North Carolina, 27708, United States; Marsico Lung Institute, University of North Carolina at Chapel Hill, Chapel Hill, North Carolina, 27599, United States; Institute for Chemical Reaction Design and Discovery, Hokkaido University, Sapporo, 001-0021, Japan

## Abstract

In the active loop extrusion model, the cohesin protein complex creates chromatin loops in eukaryotic cells. Extrusion maintains topologically associated domains (TADs), which are contiguous segments of chromatin that preferentially colocalize in space and are typically bounded by CTCF proteins that pause cohesin translocation. Here, we model active loop extrusion with hybrid molecular dynamics – Monte Carlo simulations in entangled flexible linear polymer melts. Intra-chain contact probabilities of polymers with active loop extrusion are enhanced compared to their equilibrium, passive counterparts. Extrusion causes the size of chain segments to be much smaller than in passive melts. While the overlap parameter in passive melts without extrusion monotonically increases with segment length, it is nonmonotonic in active melts and on the order of unity within the parameters of this study. Active loop extrusion suppresses contacts between TADs in favor of intra-TAD contacts. Reduction of overlaps between chain segments dilutes entanglements in active melts. Depending on parameters, active extrusion without TADs may induce more compact conformations than with TADs, due in part to fractal loopy globule-like dynamics. This work suggests that active loop extrusion reduces overlaps between TADs, contributing to effective gene regulation by *cis*-regulatory elements.

## INTRODUCTION

Active materials are out-of-equilibrium systems that convert energy from one form to another. Synthetic active materials often take inspiration from biology, which leverages myriad active processes to function and live. Biological active processes typically convert chemical energy released by ATP hydrolysis to mechanical work. In this paper, we consider a specific biological active process, called active loop extrusion, that describes how motor proteins shape the conformations of chromatin, a flexible polymer.^1–4^ Active loop extrusion is a contractile process; motor proteins pull on and compact polymer segments. Some other examples of contractile active processes are myosin motors pulling on semiflexible actin filaments and light-inducible rotary molecular motors in active gels.^5–10^

This paper is organized as follows. First, we provide biological context for active loop extrusion. We then summarize a scaling model^11^ that predicts how loop extrusion affects polymer behavior and briefly describe simulation methods used in this study. The main section of this paper presents simulation results of flexible linear polymer melts that show how contractile active extrusion compacts chains, reduces overlaps between chain segments, and suppresses entanglements.

Chromatin conformation and dynamics during interphase are crucial to gene expression patterns in eukaryotic cells, for example through increasing the probability of contacts between *cis*-regulatory elements and gene promoters.^12–14^ Both passive (thermally-driven) and active (energy-consuming) processes affect how genes are transcribed.^15–18^ Active loop extrusion involves ATP-dependent translocation of SMC protein complexes like cohesin along DNA to form progressively larger loops.^1–4^ Several studies have performed *in vitro* experiments of entangled DNA solutions with active SMC proteins from yeast.^19,20^ Cohesin is believed to actively extrude chromatin during interphase, forming topologically associated domains (TADs) demarcated by CCCTC-binding factor (CTCF) binding sites (“TAD anchors”).^21–23^ TAD anchors pause extrusion in an orientational manner; a particular anchor can only stop a cohesin domain moving in a specific direction.^21^ Genomic contacts are enriched within and suppressed between TADs.^24,25^

Previous studies developed theoretical and simulation models of active loop extrusion consistent with experimental contact maps and contact probabilities.^3,4,11,26–32^ Consider a polymer system where the passive monomeric mean square displacement (MSD) without active loop extrusion has a time dependence of ∼ Δ*t*^*α*^ where Δ*t* is the lag time. The model in ref. (11) posits that the interplay between the active kinetics of extrusion and sub-diffusive monomeric motion compacts polymers to anomalously high fractal dimensions of *D* ≈ 2/*α* on intermediate length scales. Polymer segments that have time to fully relax during the extrusion process do not compact and remain almost unperturbed. Larger sections that do not have time to equilibrate are smaller than their passive equilibrated counterparts. The model suggests that during interphase in mammalian cells, active loop extrusion compacts chromatin to a fractal dimension *D* ≈ 4 on genomic length scales between ≈ 30 kilo-base pairs (kbp) and ≈ 400 kbp, which suppresses spatial overlaps between neighboring TADs. This is consistent with genome-wide Micro-C experiments that quantify the probability that two genomic loci are in spatial proximity, and also with experiments suggesting that extrusion is important for enhancer-promoter pairs separated by more than tens of kbp.^33–35^

We extend the active loop extrusion model from ref. (11) to coarse-grain hybrid molecular dynamics – Monte Carlo (MD – MC) simulations in entangled flexible linear polymer melts. The high density of polymer melts allows for characterization of overlaps between chromatin segments as well as of entanglements (or lack thereof). Typical flexible linear polymers in melts are entangled, meaning they are topologically constrained by one another;^36^ however, experiments suggest that mammalian chromosomes do not interpenetrate significantly,^37^ and TADs tend to be spatially segregated.^38–40^ The results presented here agree with the theoretical model in ref. (11) and show that active loop extrusion suppresses inter-TAD contacts and entanglements.

## RESULTS AND DISCUSSION

### Simulation Overview

We base our simulations on the Kremer-Grest model for polymer melts.^41^ Briefly, 200 bead-spring chains of *N =* 200, 400, 800, or 1600 beads each were simulated in cubic boxes with periodic boundaries such that the bead number density was *ρ* ≈ 0.85*σ*^−3^where *σ* is the bead diameter. All polymer melts were monodisperse, and chains were fully flexible with no bending stiffness. Active loop extrusion is modeled as in ref. (11), where each cohesin is represented by an additional bond that can change binding partners. Cohesin bonds can only connect intra-chain beads. Each cohesin is modeled as two domains that each extrude with speed *v*_*ex*_. Parameters were chosen such that the expected processivity *λ* (average unimpeded loop length extruded per cohesin) and separation *d* (inverse linear number density of cohesins) were approximately equal, in line with extrusion in interphase,^4,29,42,43^ either both 25, 50, 100, or 200 beads. Most of the results in the main text use *λ* ≈ *d* ≈ 200 beads with *v*_*ex*_ ≈ 0.02 beads/*τ* unless otherwise stated, where *τ* is the Lennard-Jones time scale. Simulations with active extrusion were run without TAD anchors (denoted aNT) and with directional TAD anchors placed in convergent orientations (see Fig. S3(b)). TAD lengths were either uniform lengths of *λ* ≈ *d* throughout the simulation box (denoted aUT) or chosen from a distribution (denoted aDT) consistent with experimental Micro-C data^44^ with mean length 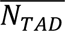 ≈ *λ* ≈ *d* unless noted otherwise. Simulations with *N* = 200 and *N* = 400 beads per chain were also run without extrusion (denoted p). See “Methods” and Supplementary Information for additional details.

### Chain compaction to anomalously high fractal dimensions

Active extrusion compacts individual chains compared to chains without extrusion. Figs. 1(a) and 1(b) show the mean square internal distance <*r*^2^(*s*)> and contact probability *P*(*s*), respectively, between two beads separated by *s* beads along the same chain. The dependence of mean square internal distances on *s* defines the fractal dimension *D* such that <*r*^2^(*s*)> ∼ *s*^2/*D*^ where ∼ indicates proportionality with a dimensional prefactor. Note that under a simple mean-field approximation, the contact probability scales as ∼ *s*^−3/*D*^.^45^ For passive systems, <*r*^2^(*s*)> ∼ *s* and *P*(*s*) ∼ *s*^−3/2^ up to the chain length, as expected for Gaussian linear chains with fractal dimension *D* ≈ 2. Systems with active extrusion using *λ* ≈ *d* ≈ 200 beads, *v*_*ex*_ ≈ 0.02 beads/*τ*, and average TAD length 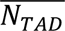 ≈ 200 beads match the passive case for small *s* because short segments have time to equilibrate during the extrusion process. We call the largest such segment the “relaxation blob” with *g*_*relax*_ beads (see Supplementary Information for additional details). For *s* > *g*_*relax*_, the active systems have an intermediate regime with shallower power law dependencies of <*r*^2^(*s*)> ∼ *s*^1/2^ and *P*(*s*) ∼ *s*^−3/4^ before returning to the *D* ≈ 2 behavior at separations of *A* = *C*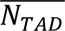, where *C* is a constant on the order of unity. The intermediate regime in these systems is indicative of activity-driven compaction, and the power laws are consistent with *D* ≈ 4. Simulations agree with genome-wide experimental contact probabilities and the theoretical model in ref. (11), where the underlying passive model is a Rouse model^36,46^ of a Gaussian polymer chain on length scales below passive entanglements.

**FIG. 1.**
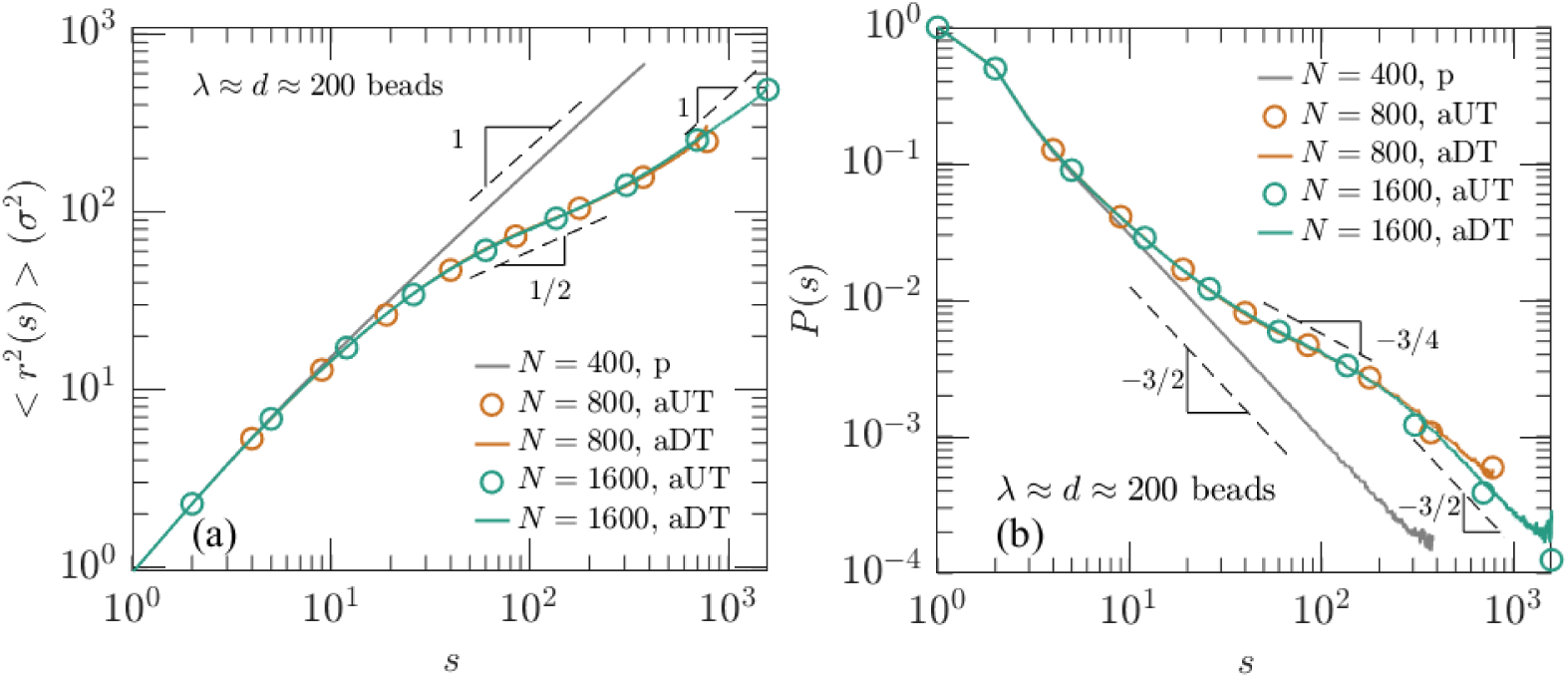
Conformations of chains in active melts with TAD anchors. (a) Mean square distances between two beads separated by *s* beads for passive melts (denoted p), active melts with uniform TAD lengths of 200 beads (denoted aUT), and active melts with a distribution of TAD lengths (denoted aDT) with average 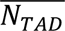 ≈ 200. Dashed lines indicate power laws. (b) Same as in (a), but for the intra-chain contact probabilities between two beads separated by *s* beads. For clarity, only ten points approximately equally spaced on the abscissa are shown for aUT melts. Orange and green solid curves almost overlap one another.

The power laws in the compact regime depend on the presence of TAD anchors and extrusion parameters. In simulations with the same *λ* ≈ *d* ≈ 200 beads as in Fig. 1 but without TAD anchors, *P*(*s*) and <*r*^2^(*s*)> suggest a compact regime with a higher fractal dimension than *D* ≈ 4, closer to *D* ≈ 7 (see Figs. 2(a) and S5). Without TAD anchors, loopy structures can form anywhere along each chain, which we propose is analogous to the fractal loopy globule (FLG) model^47^ that describes nonconcatenated entangled ring polymer melts. In the FLG model, the bead MSD scales as ∼ Δ*t*^*α*^ with *α* = 2/7 assuming full entanglement tube dilation as long as the largest loops (or rings) are on the order of or larger than the entanglement scale. The model in ref. (11) suggests that the anomalous fractal dimension resulting from active extrusion would be *D* ≈ 2/*α*, which in this case is *D* ≈ 7. Similar behavior occurs in melts with *λ* ≈ *d* ≈ 100 beads without TADs (see Figs. 2(a) and S5). However, the internal distances and contact probabilities for melts with *λ* ≈ *d* ≈ 50 beads or 25 beads without TAD anchors are consistent with *D* ≈ 4 in the compact regime (see Figs. 2(b) and S5). This suggests the entanglement onset for FLGs occurs on the order of 100 beads.

**FIG. 2:**
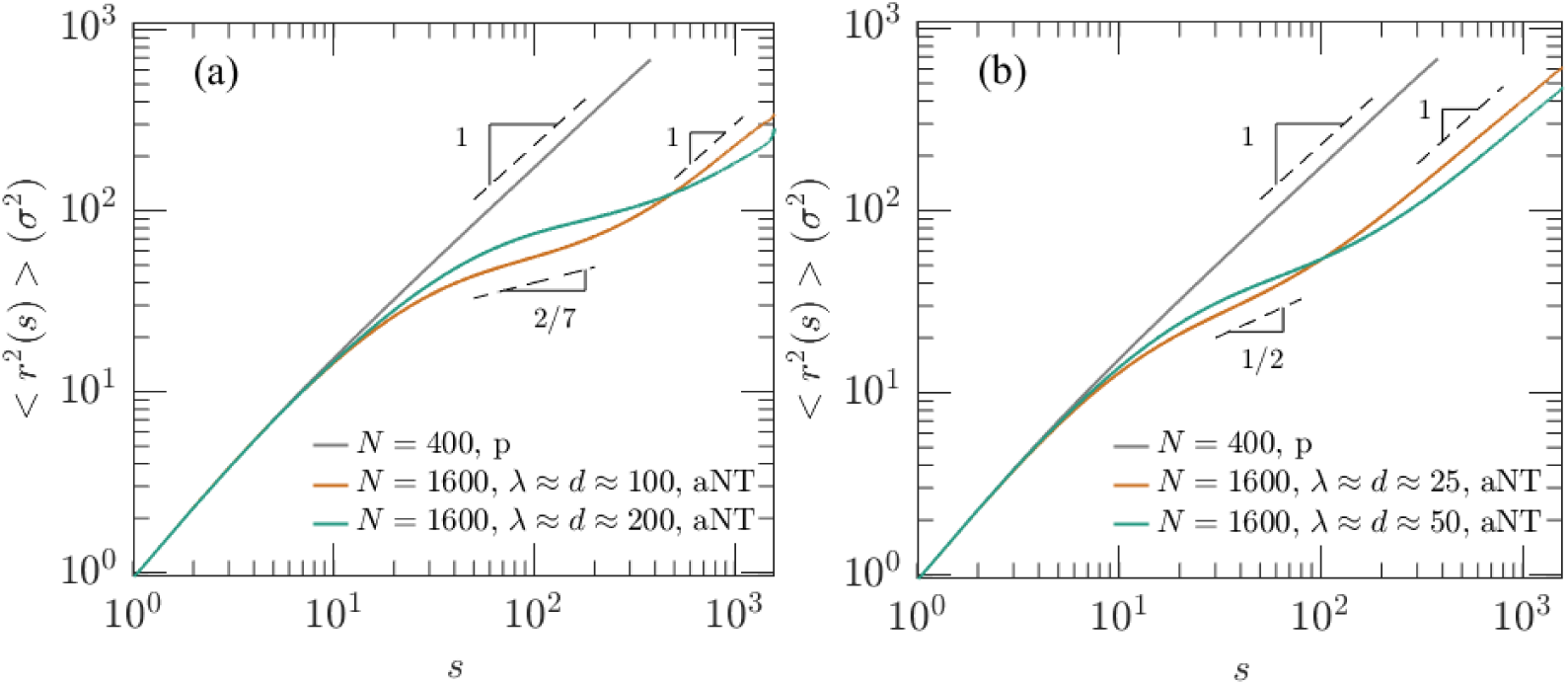
Mean square internal distances of chains in active melts without TAD anchors (denoted aNT). (a) Melts with *λ* ≈ *d* ≈100 and 200 beads. (b) Melts with *λ* ≈ *d* ≈25 and 50 beads.

With TAD anchors, loopy structures only form within predetermined boundaries with a maximum size so that each chain is a linear array of compact TADs in sequence space. The fractal structure depends on how the average TAD length 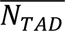 compares to processivity *λ*. In all cases, we consider approximately equal processivity and separation *λ* ≈ *d*. If 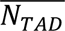 ≫ *λ*, the primary role of TAD anchors is to truncate the largest, rarest loops formed by extrusion. Chain conformations are very similar to the case without TAD anchors. On the other hand, if 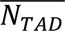 ≪ *λ*, TAD anchors can significantly suppress the effect of activity-induced compaction, because cohesins cannot extrude loops larger than TADs. Note that we consider the case where TADs are placed consecutively on each chain such that the majority of beads in the melt are encompassed within TADs (see Supplementary Information for details), similar to mammalian genomes.^48,49^ Fig. S5(b) shows that <*r*^2^(*s*)> for *λ* ≈ *d* ≈ 50 beads with 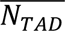 ≈ 200 beads is similar to the case of *λ* ≈ *d* ≈ 50 beads without TADs. <*r*^2^(*s*)> for *λ* ≈ *d* ≈ 200 beads with 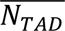 ≈ 50 is significantly larger than the case of *λ* ≈ *d* ≈ 200 without TADs. Ref. (29) suggests the boundary between these regimes is 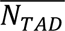 ≈ 0.6*λ*. The active melts in Fig. 1 are in the crossover regime with *λ* ≈ *d* ≈ 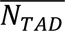. Simulations suggest that in this crossover regime, when 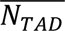 is also on the order of the FLG entanglement onset, TAD anchors suppress FLG-like behavior such that the apparent fractal dimension is *D* ≈ 4 on scales between ≈ *g*_*relax*_ and ≈ 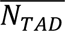. Living cells in interphase also exist in this crossover regime.

Active extrusion also changes the distribution Φ(*r*) of end-to-end distances *r* for chain segments with *s* beads (see Fig. 3(a)). The distributions for *s* < *g*_*relax*_are identical for the passive and active simulations because even with extrusion, these segments have enough time to relax. The distributions for *s* > *g*_*relax*_ are shifted to shorter distances with extrusion compared to without, even though the functional forms of the distributions are similar (see Fig. 3(b)), with subtle differences (see Fig. S6). However, the distribution of distances between TAD anchors has a significantly different form; the initial peak at short distances indicates the conformations in which TAD anchors are held together by cohesins.

**FIG. 3.**
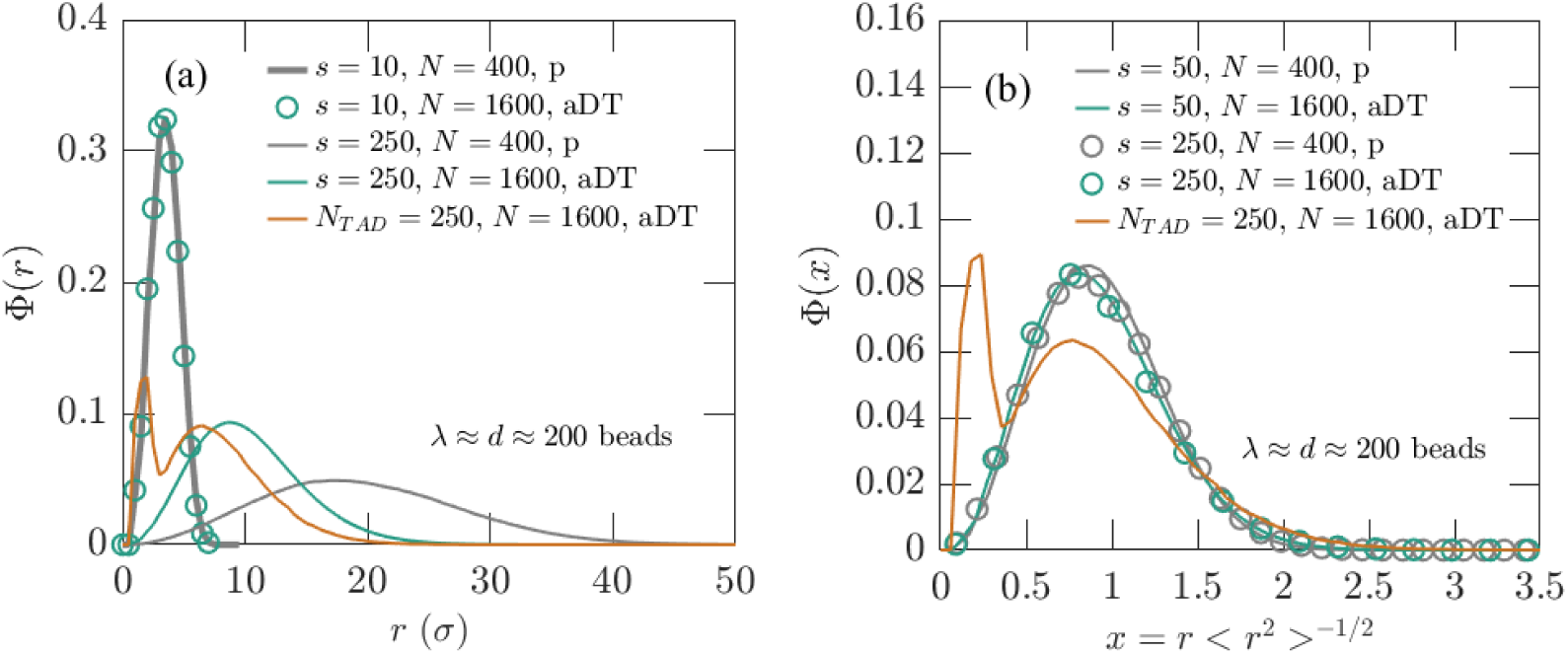
Distributions of segmental end-to-end distances. (a) Representative distributions for segments with *s* beads (gray and green) and between TAD anchors (orange). Segments with *s* = 10 beads are shorter than *g*_*relax*_while segments with *s* = 250 beads are longer. Φ(*r*) is normalized such that ∫ Φ(*r*)*dr* = 1. (b) Normalized distributions of end-to-end distances for chain segments and between TAD anchors. The abscissa is the distance divided by the root-mean-square end-to-end distance for a given segment. Φ(*x*) is normalized such that ∫ Φ(*x*)4*πx*^2^*dx* = 1.

### Nonmonotonic overlap parameter in melts with active extrusion

Active extrusion alters spatial overlaps between neighboring chain segments. Consider a segment with *s* beads and root-mean-squared radius of gyration *R*_*g*_(*s*). The overlap parameter is calculated as *O*(*s*) = *n*_*Rg*_(*s*)/*n*_*self*_(*s*), where *n*_*Rg*_(*s*) is the total number of beads within *R*_*g*_(*s*) of a segment’s center of mass (COM) and *n*_*self*_(*s*) is the number of beads belonging to the segment itself within *R*_*g*_(*s*) of the COM (see Fig. 4(a)). For systems with active extrusion, a chosen segment with *s* beads can span multiple TADs.

**FIG. 4.**
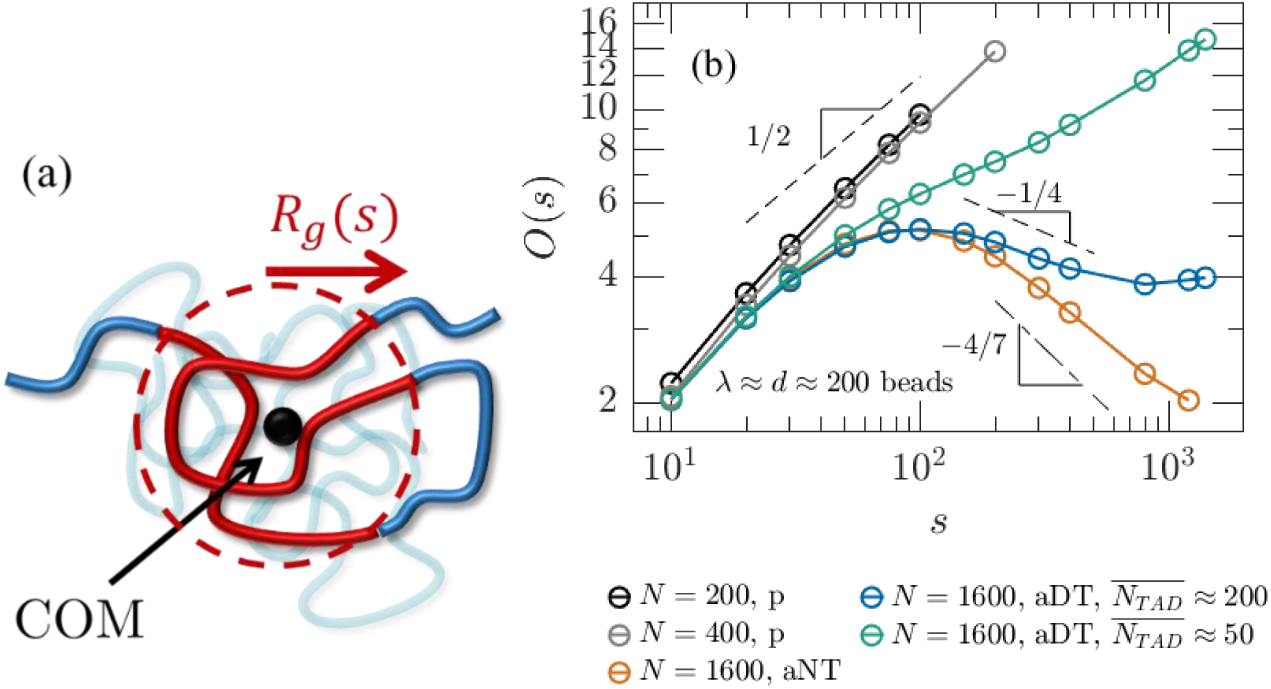
Overlap parameter in active and passive melts. (a) Schematic for calculating the overlap parameter. The thick curve (blue and red together) indicates the segment of interest with length *s* and COM at the filled black circle. The radius of the dashed red circle is the segment’s radius of gyration. The red portion of the segment contributes to *n*_*self*_. All polymer mass within the dashed red circle contributes to *n*_*Rg*_. (b) Overlap parameter for passive melts and active melts with *λ* ≈ *d* ≈ 200 beads.

As seen in Fig. 4(b), the overlap parameter *O*(*s*) in passive melts monotonically increases with segment length *s*, consistent with the scaling prediction *O*(*s*) ∼ *s*^1/2^. In contrast, in active melts with compact regimes of *D* ≈ 4 or *D* ≈ 7, the overlap parameter *O*(*s*) is nonmonotonic (representative curves shown in blue and orange, respectively). For intermediate *s*, the number of overlapping segments within a segment’s pervaded volume decreases with *s*. This is due to the anomalously high fractal dimension related to extrusion-mediated compaction; simulations are consistent with the scaling prediction^11^ of ∼ *s*^3/*D*−1^ in the region with decreasing *O*(*s*) (∼ *s*^−1/4^ for *D* ≈ 4 and ∼ *s*^−4/7^for *D* ≈ 7).

The precise shape of *O*(*s*) depends on extrusion parameters and the presence of TADs, as they dictate the fractal structure in the regime of activity-induced compaction (see Figs. S1 and S7 and discussion following Fig. 1). For example, when 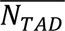 < *λ*, TAD anchors restrict extrusion-associated compaction resulting in a monotonically increasing *O*(*s*) (see green curve in Fig. 4(b)).

For parameters corresponding to a nonmonotonic *O*(*s*), ref. (11) predicts the local maximum to be at contour lengths on the order of the relaxation blob. The local minimum is predicted to be on the order of the average TAD length or cohesin processivity with or without TADs, respectively. Figs. 4(b) and S8 suggest that the maximum occurs at ≈ 5*g*_*relax*_; with TADs and *λ* ≈ *d* ≈ 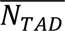, the minimum occurs at ≈ 4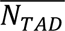; without TADs or if 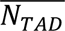 > *λ* ≈ *d*, the local minimum occurs at ≈ 8*λ*. Even if the decreasing portion of *O*(*s*) spans almost a full decade in contour length, the weak power law means that the overlap parameter varies by much less than a decade in the same regime. The shape of *O*(*s*) remains similar but vertically shifted to smaller values (fewer overlaps) if a chain segment’s pervaded volume is considered to be an effective ellipsoid defined by its gyration tensor rather than a sphere (see Fig. S9(c)).

### Ellipsoidal radial distribution functions of TADs

The volume explored by a TAD is enriched with its own mass. We characterize this by calculating “ellipsoidal” radial distribution functions (eRDF) of TADs measured from their COMs. The eRDF is an ellipsoidal analogue of a conventional radial distribution function. Since TADs are not spherical (see Fig. S9(a)), we consider the pervaded volume of a TAD to be an ellipsoid with axes defined by its gyration tensor. The value of eRDF at *r*_*e*_ reflects the density of beads within an ellipsoidal shell with inner radii *r*_*e*_*λ*_1_, *r*_*e*_*λ*_2_, and *r*_*e*_*λ*_3_ and outer radii (*r*_*e*_ + Δ*r*_*e*_)*λ*_1_, (*r*_*e*_ + Δ*r*_*e*_)*λ*_2_, and (*r*_*e*_ + Δ*r*_*e*_)*λ*_3_, where *λ*_1_, *λ*_2_, and *λ*_3_ are the square roots of the gyration tensor eigenvalues (see Supplementary Information for additional details). We decompose the eRDF into the “self” (eRDF_s_) and “other” (eRDF_o_) components, which reflect the densities of beads belonging to a TAD of interest itself or coming from other TADs, respectively.

Fig. 5(a) shows eRDF_s_ and eRDF_o_ for simulations with *λ* ≈ *d* ≈ 200 beads and a distribution of TAD lengths. Within the ellipsoidal pervaded volume of a given TAD (*r*_*e*_ ≤ 1), eRDF_s_ is greater than eRDF_o_ for most TAD lengths, and both are relatively flat, indicating an enrichment of “owned” beads belonging to this TAD with nearly constant density. eRDF_s_ for *r*_*e*_ ≤ 1 increases with TAD length. Longer TADs also promote TAD segregation. Increasing TAD length narrows the peak of the product eRDF_s_∗eRDF_o_ (see Fig. 5(b)), suggesting a transition to a TAD morphology with a center nearly free of beads from other TADs and an interpenetration zone at the interface with other TADs, reminiscent of polymer-grafted nanoparticle (GNP) melts.^50^ In contrast to GNP melts, the center of TAD-pervaded volumes is not completely free of beads from other TADs. Fig. S9(b) plots *r*_*e*,1/2_, which is where eRDF_s_ = eRDF_o_ = 0.5, indicating where the majority component switches from beads owned by the TAD of interest to beads from other TADs. *r*_*e*,1/2_ ≈ 1.5 for our longest TADs of *N*_*TAD*_ = 800 beads, suggesting that these long TADs can occupy the majority of space approximately three times the ellipsoidal volumes defined by their gyration tensors. This implies that their actual pervaded volumes are significantly larger.

**FIG. 5.**
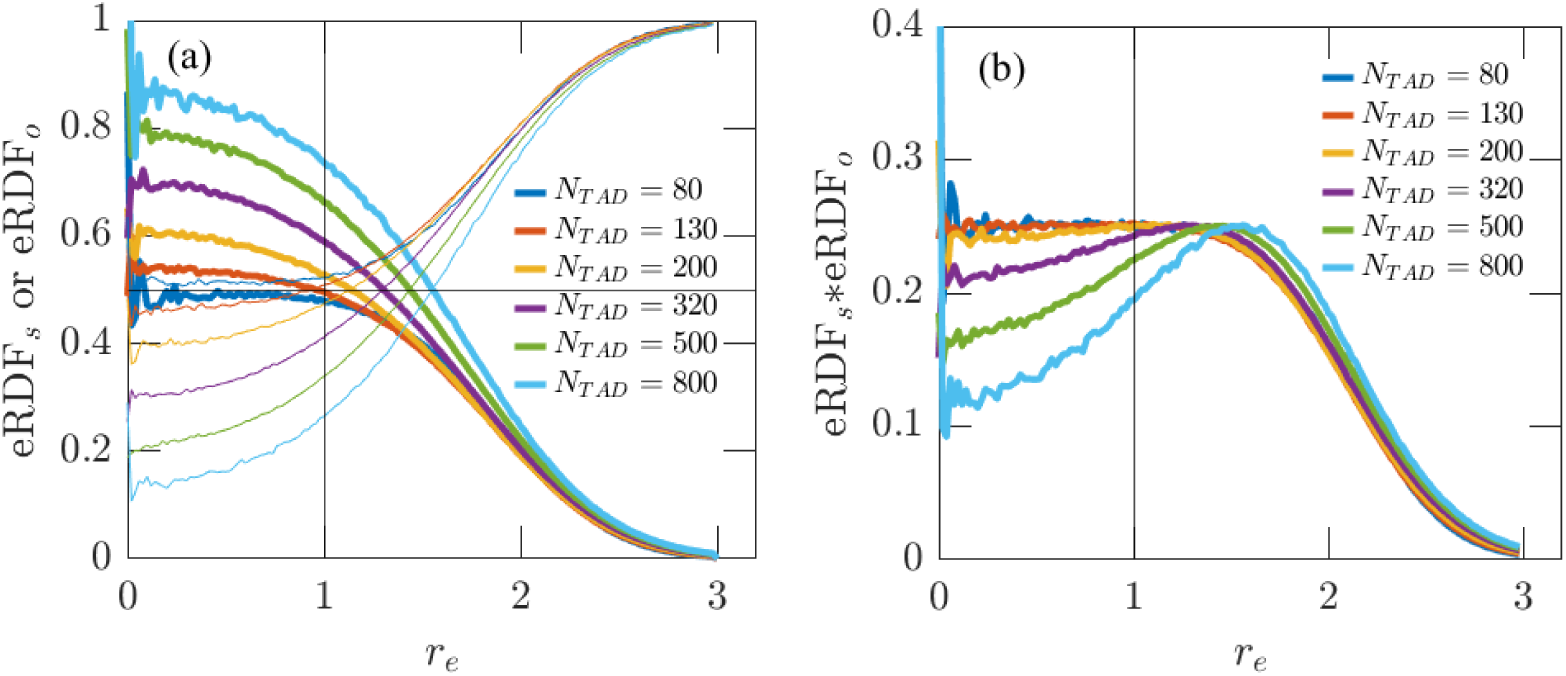
Ellipsoidal radial distribution functions (eRDFs) of active melts with TADs. (a) eRDFs measured from TAD COMs in active melts with *N* = 1600 beads per chain, *λ* ≈ *d* ≈ 200 beads, and a distribution of TAD lengths throughout the melt. The “self” component eRDF_s_ and “other” component eRDF_o_ are shown in thick and thin curves, respectively. Vertical line at *r*_*e*_ = 1 corresponds to the normalized “radius” of ellipsoidal pervaded volumes. Horizontal line at 0.5 corresponds to equal contributions from the self and other components. (b) eRDF_s_∗eRDF_o_ for the active melts shown in (a), suggesting the development of an interpenetration zone at the interface between TADs for larger *N*_*TAD*_.

### Enhancement of intra-TAD contacts

Overlap suppression due to active loop extrusion suggests that the fraction of intra-TAD contacts made by loci within a given TAD is an increasing function of TAD length. Intra-TAD contacts are those made by beads of a TAD with other beads within the same TAD. Here, we only consider intra-chain contacts (or intra-chromosomal contacts in the case of experiments). In simulations, inter-TAD contacts are between monomers of different TADs along the same chain. Fig. 6 shows the intra- and inter-TAD contact fractions for two cell types (HFFs^44^ and mESCs^51^) and active melt simulations with *λ* ≈ *d* ≈ 200 beads and a distribution of TAD lengths with 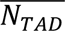 ≈ 200 beads. We also show theoretical predictions based on genome- or melt-wide *P*(*s*) curves. The fraction of intra-TAD contacts *F*_*intra*_(*N*_*TAD*_) is estimated as

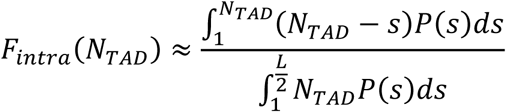

**FIG. 6.**
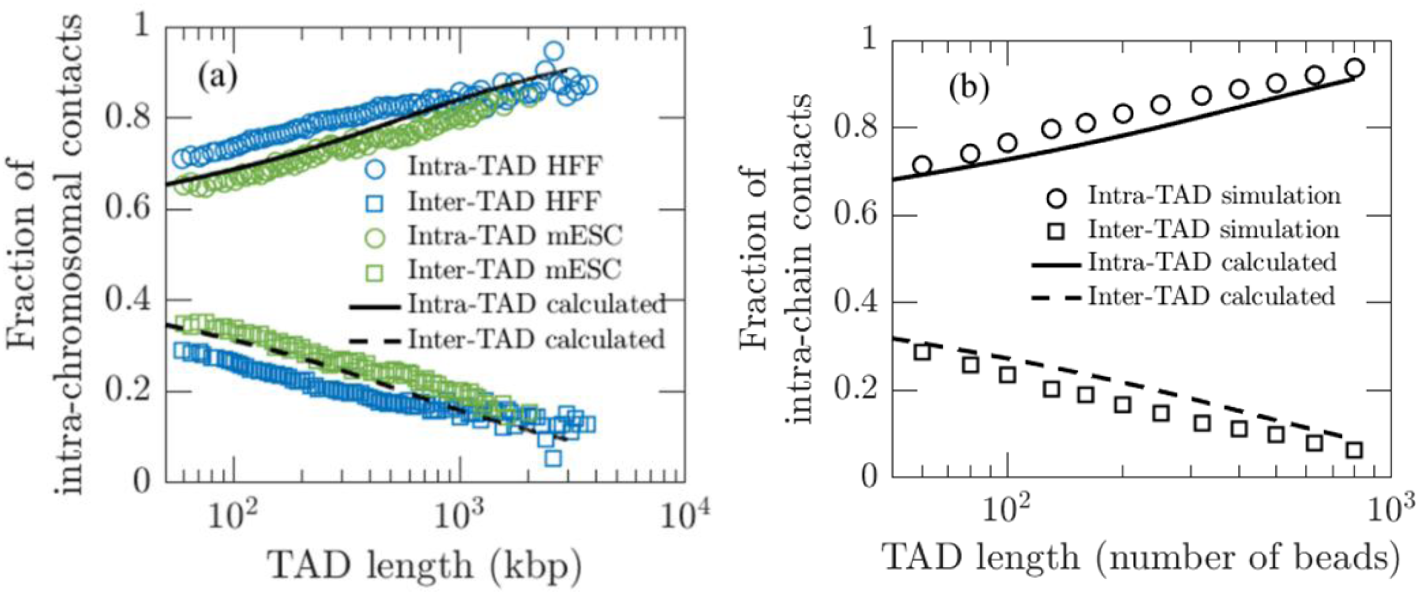
Intra-and inter-TAD contact fractions. (a) Intra- and inter-TAD contact fractions of all intra-chromosomal contacts from experimental Micro-C in HFF^44^ and mESCs^51^. Calculated predictions used *g*_*relax*_ = 26 kbp, *A* = 350 kbp, and *L* = 100 megabase pairs. (b) Intra- and inter-TAD contact fractions of all intra-chain contacts in active melt simulations with *N* = 1600 beads per chain, *λ* ≈ *d* ≈ 200 beads, and a distribution of TAD lengths throughout the melt. Calculated predictions used *g*_*relax*_ = 15 beads, *A* = 268 beads, and *L* = 1600 beads.

(Eq. S8 in the Supplementary Information), where the numerator is approximately the number of intra-TAD contacts, the denominator is approximately the total number of intra-chain contacts, *L* is the average chain (or chromosome) length, *N*_*TAD*_is the TAD length, and *P*(*s*) is the contact probability function. For the cases considered here, *P*(*s*) scales as ∼ *s*^−3/2^ for *s* < *g*_*relax*_, ∼ *s*^−3/4^ for *g*_*relax*_ ≤ *s* < *A*, and ∼ *s*^−3/2^ for *A* ≤ *s*, where *g*_*relax*_ is the relaxation blob length and *A* = *C*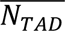 where *C* is a constant on the order of unity. The intra-TAD contact fraction is then approximately

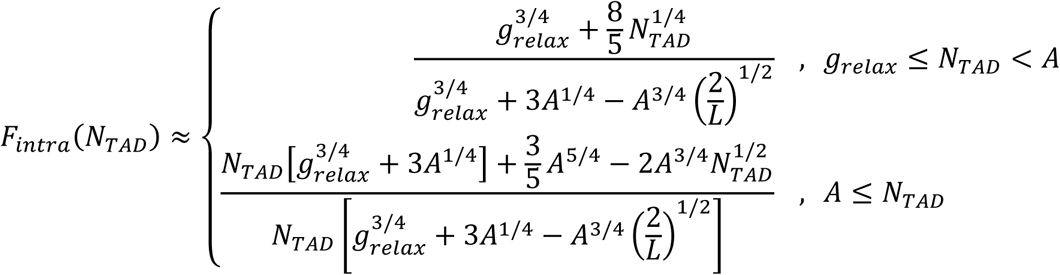

See section I.d. and Eqs. S6 – S12 in the Supplementary Information for details. The theoretical predictions for the intra-TAD contact fraction *F*_*intra*_(*N*_*TAD*_) (solid black curves) and the inter-TAD contact fraction *F*_*inter*_(*N*_*TAD*_) = 1 − *F*_*intra*_(*N*_*TAD*_) (dashed black curves) have good agreement with experimental and simulation results.

### Entanglement dilution in active melts

Another consequence of extrusion-mediated overlap reduction is entanglement dilution. Using the Z1+ package,^52^ we find that the apparent entanglement strand *N*_*e*_in melts with active loop extrusion is significantly larger than in the passive melt counterparts (see Fig. 7 and Supplementary Information for details). Note that existing algorithms for quantifying entanglements may not perform well on weakly entangled systems.^53^ Without TAD anchors, the apparent *N*_*e*_ increases almost linearly with chain length, with one to three apparent entanglements per chain (see Fig. S10(c)), suggesting that chains are barely entangled or just at the crossover to entanglements, regardless of chain length. This is analogous to FLGs, where ring polymers adopt conformations that are close to the entanglement threshold on all length scales.^47^ The overlap parameter varies very little (see Figs. 4(b) and S10), reminiscent of the constant overlap parameter in FLGs. It is also possible that chains need to be much longer to measure true entanglements of these systems.

**FIG. 7.**
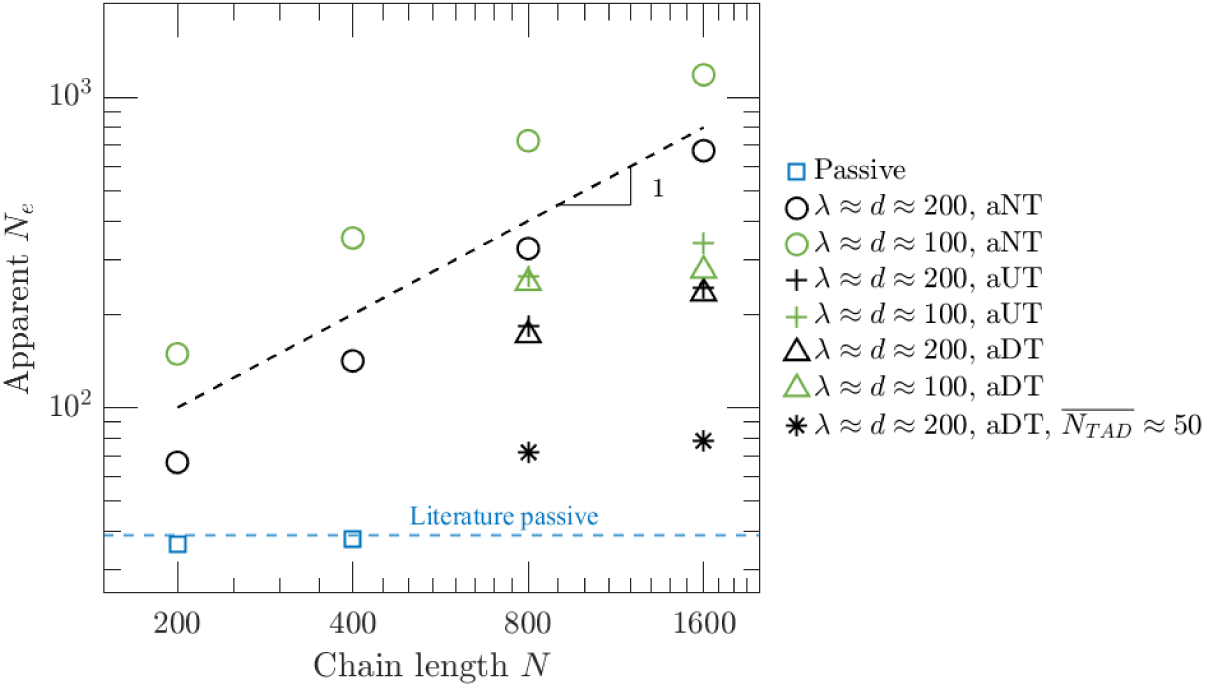
Number of beads in an apparent entanglement strand *N*_*e*_ calculated with Z1+.^52^ Active melts with TADs shown here have *λ* ≈ *d* ≈ 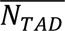 except for black stars.

Simulations with TAD anchors and *λ* ≈ *d* ≈ 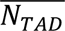 have a smaller apparent *N*_*e*_ than without anchors, though still much larger than the passive case. This reflects that the fractal dimension without TAD anchors is much larger than with TAD anchors (*D* ≈ 7 versus *D* ≈ 4). For these simulations with *λ* ≈ *d* ≈ 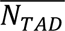, the overlap parameter at the apparent *N*_*e*_ is close to four, approximately the same as the overlap parameter at the passive *N*_*e*_. This is consistent with the prediction that the same number of chain sections overlap at *N*_*e*_ for both active and passive cases (in ref. (11), this overlap parameter was assumed to be ten). We note that the theory assumed that entanglements can be determined by a characteristic overlap parameter calculated using spherical pervaded volumes of chain segments in both active and passive cases, which is likely an oversimplification. For the longest chains studied with *λ* ≈ *d* ≈ 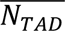, the ratio of the apparent *N*_*e*_ to 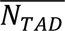 is between 1 and 4 depending on *λ* (see triangles and plus signs in Figs. 7 and S10). As such, TADs shorter than 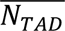 are not entangled, while TADs much longer than 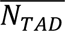 can entangle. In other words, shorter TADs are not significantly restricted by spatially neighboring TADs. The ratio *N*_*e*_/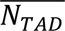 increases if *λ* < 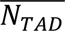 (see Fig. S10) because chains behave similarly to the case without TAD anchors.

Simulations with 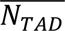 < *λ* ≈ *d* still have larger *N*_*e*_ than the passive case but smaller than when 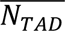 ≈ *λ* ≈ *d*, reflecting that activity-induced compaction is restricted in these conditions (see stars in Fig. 7). Fig. S10 shows the apparent *N*_*e*_ for several other simulation conditions. Reduction of entanglements was previously shown in simulations of active loop extrusion in semiflexible polymer solutions as well as passive melts of flexible loopy polymers.^27,54^ Our work demonstrates this effect in dense, flexible polymer melts with active extrusion.

### Summary

In summary, we present simulations of active loop extrusion in flexible linear polymer melts and find agreement with scaling model predictions.^11^ Active extrusion enhances intra-chain contact probabilities *P*(*s*) compared to passive melt counterparts and can compact polymers to anomalously high fractal dimensions of *D* ≈ 4 (Figs. 1 and 3) or *D* ≈ 7 (Figs. 2 and S5) depending on extrusion parameters and the presence of TAD anchors. Active extrusion without TAD anchors may induce FLG-like conformations if cohesin processivity is larger than the entanglement scale, consistent with the almost linear dependence of apparent *N*_*e*_ on chain length and a weakly varying overlap parameter over a wide range of contour lengths (see Figs. 4, 7, S7, S8, and S10). The relative values of average TAD length and extrusion processivity also determines whether or not extruder activity effectively compacts chains. This work shows that active loop extrusion causes a nonmonotonic dependence of the overlap parameter *O*(*s*) on segment length *s* and suppresses entanglements (Figs. 4, 7, S7, S8, and S10). The pervaded volume of a given TAD is enriched with its own mass compared to mass from other TADs, and simulations suggest that longer TADs develop an interpenetration zone at the interface between TADs (Fig. 5). The fraction of intra-TAD contacts in our simulations and experimental data agrees with theoretical predictions (Fig. 6). Our work supports the hypothesis that active loop extrusion compacts TADs, segregates them, and enhances intra-TAD contacts to facilitate efficient transcriptional regulation.

## METHODS

Hybrid MD – MC simulations are based on the method in ref. (11). Extrusion was implemented as a custom “fix” in the LAMMPS simulation package.^55,56^ Cohesins bind to random pairs of neighboring beads on the same chain with probability 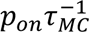 per pair of beads and randomly unbind with probability 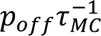 per bound cohesin (where *τ*_*MC*_ is the Monte Carlo timestep) such that there is an average number density of bound cohesins in steady state of either one cohesin per *p*_*off*_/*p*_*on*_ =25, 50, 100, or 200 beads. Cohesins are modeled as two domains that each extrude at an average speed of one bead per 25 *τ* or 50 *τ* (*v*_*ex*_ ≈ 0.04 beads/*τ* or *v*_*ex*_ ≈ 0.02 beads/*τ*, respectively) regardless of the presence of other cohesins (unimpeded cohesin traversal). We define two beads in our simulations to be in contact if the distance between their centers of mass is at most 1.5*σ*. Most simulations were run for more than ten times the end-to-end vector autocorrelation decay time. See Supplementary Information for additional simulation details. Tables S1 and S2 show the parameter combinations used in this work.

## ACKNOWLEDGEMENTS

This work was performed, in part, at the Center for Integrated Nanotechnologies, an Office of Science User Facility operated for the U.S. Department of Energy (DOE) Office of Science by Los Alamos National Laboratory (Contract 89233218CNA000001) and Sandia National Laboratories (Contract DE-NA-0003525). The authors thank Dr. Gary Grest for access to initial chain coordinates for simulations.

## FUNDING SOURCES

The authors acknowledge support by the NIH under grant number P01HL164320 and the NSF under grant number 2621977.

## AUTHOR CONTRIBUTIONS

BC and MR designed the research; BC performed the research; BC and MR analyzed the data; BC and MR wrote the paper.

## COMPETING INTERESTS

The authors declare no competing interests.

## Notes

### Competing Interest Statement

The authors have declared no competing interest.

