## Supplemental Information for "Entanglement dilution and high fractal dimension mediated by loop extrusion revealed in simulations of active polymer melts"

**Table of contents:**

I. **Extended methods**
   a. Hybrid molecular dynamics – Monte Carlo simulations
   b. TAD length distributions
   c. Ellipsoidal pervaded volumes and ellipsoidal radial distribution functions (eRDF)
   d. Intra- and inter- TAD contact fractions
   e. Apparent entanglement strand length

II. **Extended data**
   a. Internal distances and contact probabilities
   b. Segment size distributions
   c. Overlap parameters
   d. Anisotropy, $r_{e,1/2}$, and ellipsoidal overlap parameter
   e. Apparent entanglement strand

### I. Extended methods

#### a. Hybrid molecular dynamics – Monte Carlo (MD—MC) simulations

Simulations of monodisperse polymer melts were performed based on the Kremer-Grest bead-spring model.[1] Simulations used the Large-scale Atomic/Molecular Massively Parallel Simulator (LAMMPS) package.[2] Each simulation had 200 chains, each with $N = 200$, 400, 800, or 1600 beads of diameter $\sigma$. Non-bonded beads interact with a shifted and truncated Lennard-Jones (LJ) potential:

$$U_{LJ}(r) = \begin{cases} 4\epsilon_{LJ}\left[\left(\frac{\sigma}{r}\right)^{12} - \left(\frac{\sigma}{r}\right)^{6} - \left(\frac{\sigma}{r_c}\right)^{12} + \left(\frac{\sigma}{r_c}\right)^{6}\right] & , r \leq r_c \\ 0 & , r > r_c \end{cases} \tag{S1}$$

with the distance cut-off $r_c = 2^{1/6}\sigma$ and the LJ energy $\epsilon_{LJ}$. The mass of a bead is $m$, and the LJ time is $\tau = \left(m\sigma/\epsilon_{LJ}\right)^{1/2}$. Bonds along the chain backbone are represented by a finite extensible nonlinear elastic (FENE) potential:

$$U_B(r) = -0.5KR_0^2 ln\left[1 - \left(\frac{r}{R_0}\right)^2\right] + U_{LJ}(r)\,, \tag{S2}$$

where $K = 30\epsilon_{LJ}\sigma^{-2}$, $R_0 = 1.5\sigma$, and $U_{LJ}(r)$ is given by Eq. (S1). We start with nonconcatenated semiflexible rings with an angle potential $U_{angle} = 1.5\epsilon_{LJ}[1 + cos(\theta)]$ in a large cubic box that was compressed to a target pressure of $P = 5\epsilon_{LJ}/\sigma^3$ in the absence of activity using NPT simulations, resulting in a bead number density of $\rho \approx 0.85$ beads/$\sigma^3$. During the NPT run, temperature was maintained at $T = 1\epsilon_{LJ}/k_BT$, the damping constant for both thermostat and barostat was $0.01\tau^{-1}$, and the integration timestep was $\Delta t = 0.01\tau$. After $10^4\tau$ of NPT, the rings were cut once to linearize them and the angle potential was turned off. Simulations were switched to NVT ensembles where a Langevin thermostat was used to maintain a temperature of $T = 1\epsilon_{LJ}/k_BT$ with a damping constant $\Gamma = 1\tau^{-1}$. Activity was then turned on and modeled via the method described below. Passive melts with $N = 200$ or $N = 400$ beads per chain were run for $> 3 * 10^6\tau$ and $> 7 * 10^6\tau$ in the absence of activity, respectively.

We chose to prepare our simulations with linearized ring melts so that the initial chain conformations are relatively compact compared to relaxed linear melts, which reduces the time needed for long-chain melts to reach a steady-state with activity. For a melt with $N = 200$ beads per chain without TADs and $\lambda \approx d \approx 200$ , we also ran an active extrusion simulation using an initial conformation from the end of the passive run. We did not observe differences between mean squared internal distances $<r^2(s)>$ of active melts using the different initial conformations (see Fig. S1(a)), which we believe justifies our choice of initial conformations for active melts with longer chains. Additionally, for a melt with $N = 800$ beads per chain without TADs and $\lambda \approx d \approx 50$, we ran an active extrusion simulation using an initial conformation of beads placed on a regular lattice at a number density of $\rho \approx 0.85$ beads/$\sigma^3$. We did not observe differences in $<r^2(s)>$ between the initial conformation of a regular lattice compared to linearized rings (see Fig. S1(b)). Having shown that the initial conformation does not impact the steady-state conformation of our active melts, we randomly chose conformations from the trajectories of some active melts as initial conformations of other active melts.

In most cases, the time interval used to collect data was more than four times the end-to-end vector autocorrelation time (see Fig. S1(c)), and in many cases more than ten times. During this time, the mean square displacement (MSD) of a bead ranged between 1 and 10 times the mean squared end-to-end distance of a chain. The only case where the simulation was not run for this long was the melt with $N = 1600$ beads, $\lambda \approx d \approx 200$, and a distribution of TADs with $\overline{N_{TAD}} \approx 50$. This is the case where TAD anchors strongly suppress activity-induced compaction. Even after $\approx 1.4 * 10^7\tau$, the autocorrelation remained larger than 0.4. For this melt, data were collected for the second half of the simulation run. The chain statistics in this system were essentially the same as for melts with $N = 800$ beads, $\lambda \approx d \approx 200$, and a distribution of TADs with $\overline{N_{TAD}} \approx 50$ (see Figs. S5(b) and S5(d)).

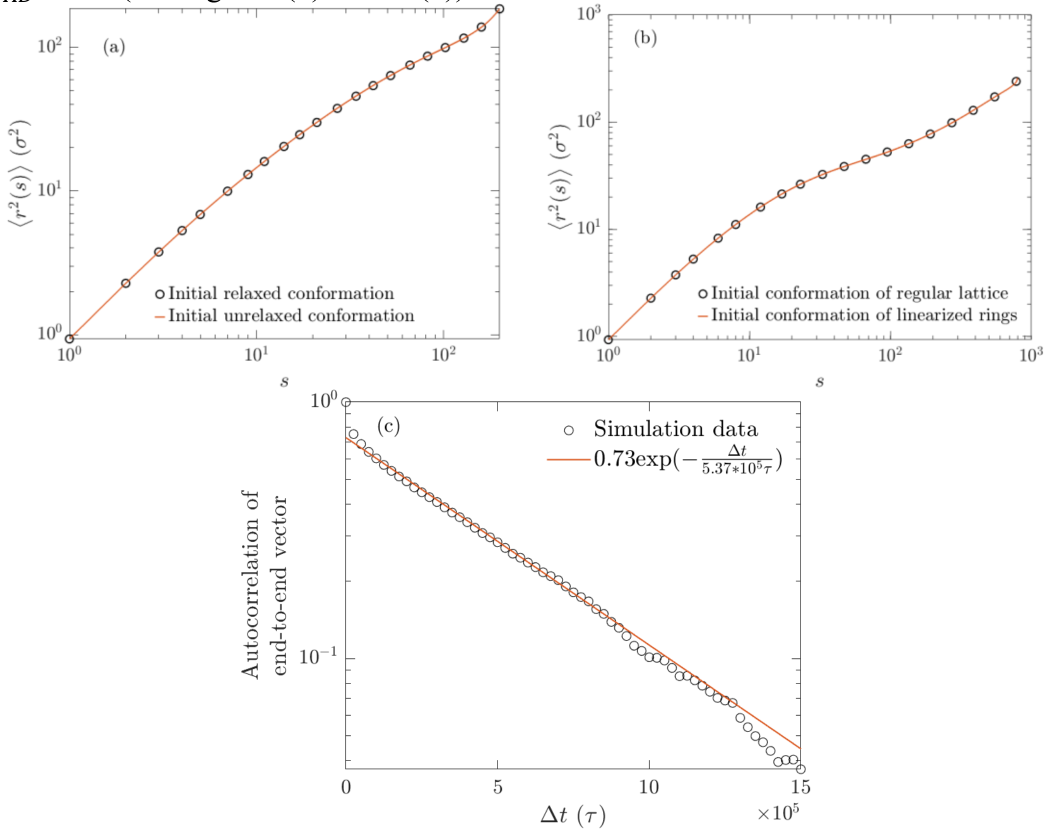


**Figure S1**: (a) Comparison of mean square internal distances of active melts with $N = 200$ beads and $\lambda \approx d \approx 200$ without TADs with different initial conformations. For clarity, only 21 uniformly-spaced points along the logarithmic axis are shown for the melts with initially relaxed chains (black circles). (b) Comparison of mean square internal distances of active melts with $N = 800$ beads and $\lambda \approx d \approx 50$ without TADs with different initial conformations. For clarity, only 19 uniformly-spaced points along the logarithmic axis are shown for the melt that started from an initial lattice conformation (black circles). (c) Example autocorrelation of the end-to-end vector of chains in a melt with $N = 1600$ beads per chain and $\lambda \approx d \approx 50$ without TADs. The red curve is a fit to lag times between $\Delta t = 2.5 * 10^5\tau$ and $\Delta t = 10^6\tau$. The decay time is approximately $(5.37 \pm 0.07) * 10^5\tau$.

Active extrusion was modeled with hybrid MD—MC as described previously[3] where MD was used to follow bead trajectories, and MC was used to determine cohesin/extruder binding, unbinding, and translocation. The MC method for active extrusion was implemented as a custom "fix" in LAMMPS.[11] MC steps were performed every $\tau_{MC}$. We provide a brief description here and refer readers to ref. (3) for additional details. Briefly, cohesins/extruders are modeled as an additional bond with potential

$$U_c(r) = -0.5K_c R_{0_c}^2 ln\left[1 - \left(\frac{r}{R_{0_c}}\right)^2\right] + U_{LJ}(r)\,, \qquad \text{(S3)}$$

where $K_c = 1\epsilon_{LJ}\sigma^{-2}$, $R_{0_c} = 4\sigma$, and $U_{LJ}(r)$ is given by Eq. (S1). This bond can only act on intra-chain beads, and can be added, broken, or moved during MC steps. During each MC step, the order of cohesin binding, unbinding, and translocation is randomized. The cohesin bond is added to a random pair of neighboring intra-chain beads $x$ and $x + 1$ with probability $p_{on}$ per $\tau_{MC}$. During each MC step, all possible pairs of neighboring intra-chain beads are cycled through in random order for potential binding. Each bound cohesin bond breaks with probability $p_{off}$ per $\tau_{MC}$.

For translocation, each domain or "side" of the cohesin bond attempts a move to a neighboring bead (e.g., from $x$ to $x - 1$ and from $x + 1$ to $x + 2$) with probability $p = 0.5$ such that the theoretical average extrusion rate per domain is $v_{ex} = 0.5$ beads/$\tau_{MC}$ and the theoretical average *total* extrusion rate is $2v_{ex} = 1$ bead/$\tau_{MC}$. The choices of $\tau_{MC}$, $v_{ex}$, $p_{off}$, and $p_{on}$ define the theoretical cohesin processivity $\lambda$ and separation $d$ where $\lambda = 2v_{ex}\tau_{MC}/p_{off} = 1/p_{off}$ beads and $d = p_{off}/p_{on}$ beads. We model unimpeded cohesin traversal, meaning that each bead can participate in more than one cohesin bond. Translocation moves are accepted unless a cohesin domain reaches a TAD anchor or the attempted bond length is longer than $R_{0_c} - 0.005\sigma = 3.995\sigma$ (to prevent over-stretched FENE bonds). If a domain attempts to move off a chain end, the cohesin unbinds. For these reasons, the actual extrusion rates, processivity, and separation are not exactly the theoretical values. We report the theoretical values for consistency. Table S1 shows the parameters used in this study.

**Table S1**: Parameters used to simulate active melts

| $\tau_{MC}$ ($\tau$) | Theoretical $v_{ex}$ (beads/$\tau_{MC}$) | Theoretical $v_{ex}$ (beads/$\tau$) | $p_{on}$ | $p_{off}$ | Theoretical $\lambda$ (beads) | Theoretical $d$ (beads) | $g_{relax}$ (beads) |
|---|---|---|---|---|---|---|---|
| 25 | 0.5 | 0.02 | $2.5\text{x}10^{-5}$ | $5\text{x}10^{-3}$ | 200 | 200 | 20 |
| 25 | 0.5 | 0.02 | $4\text{x}10^{-4}$ | $2\text{x}10^{-2}$ | 50 | 50 | 11 |
| 25 | 0.5 | 0.02 | $1.6\text{x}10^{-3}$ | $4\text{x}10^{-2}$ | 25 | 25 | 9 |
| 12.5 | 0.5 | 0.04 | $1\text{x}10^{-4}$ | $1\text{x}10^{-2}$ | 100 | 100 | 11 |

The time between cohesin binding events within a segment of length $\lambda$ is $\tau_{B,\lambda} \approx d/(2v_{ex})$. This time in part determines the length of the largest relaxation blob $g_{relax}$ reported in the last column of Table S1.[3] The parameters for simulations with $\lambda = d = 50$ beads and $\lambda = d = 100$ beads were chosen to have the same $\tau_{B,\lambda} \approx 1250\tau$ and thus the same expected largest $g_{relax}$. The parameters for simulations with $\lambda = d =$ 200, 50, and 25 beads were chosen to have the same $v_{ex}$. The number of beads in the largest relaxation blob $g_{relax}$ was calculated by a similar method to ref. (3). We compare the mean square displacement of a bead in a *passive* melt $<MSD_{p,b}>$ with $N = 400$ beads per chain to the mean square internal distances in the same *passive* melt $<r_p^2>$. $g_{relax}$ is the chain section with passive diffusion time equal to $\tau_{B,\lambda}$. We estimate $g_{relax}$ as the longest section with passive mean squared size less than or equal to the bead MSD during $\tau_{B,\lambda}$, or $<r_p^2> \leq <MSD_{p,b}(\tau_{B,\lambda})>$ (see Fig. S2).

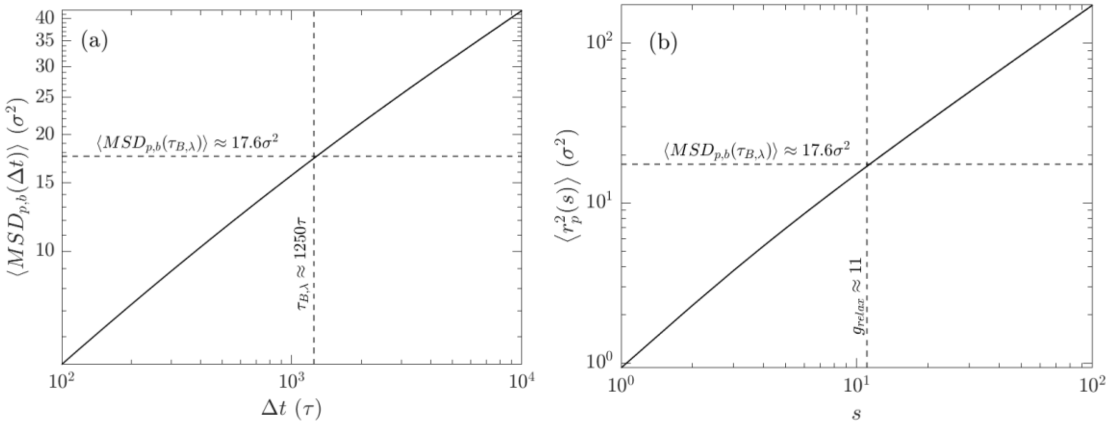


**Figure S2**: Example for determining $g_{relax}$ for $\tau_{B,\lambda} \approx 1250\tau$. a) Mean square displacement of a bead in a passive melt. Dashed vertical line shows $\tau_{B,\lambda}$, which is the expected time between cohesin binding events within $\lambda$. Dashed horizontal line shows the approximate MSD at lag time of $\tau_{B,\lambda}$. b) Mean square internal distances of a passive melt. Dashed vertical line shows the number of beads $g_{relax}$ where its passive size is approximately the passive bead MSD at $\tau_{B,\lambda}$.

#### b. TAD length distributions

Simulations of all chain lengths were performed without TAD anchors. Simulations with $N = 800$ and 1600 beads per chain were also performed with TAD anchors. Simulations with uniform TAD lengths had length $N_{TAD} = \lambda \approx d$. These TAD anchors were placed such that each chain had full coverage of TADs. For example, a chain with $N = 800$ beads and TADs with length $N_{TAD} = 200$ beads had TAD anchors placed at beads (1, 100), (101, 200), (301, 400), (401, 600), and (601, 800). Each chain in the melt with uniform TAD lengths had the same TAD anchors. TAD anchors are directional. For example, the anchor at bead index 101 only stops cohesin domains that move from bead index 102 to 101, while the anchor at index 200 only stops domains moving from index 199 to 200.

Non-uniform TAD lengths were chosen from a distribution consistent with Micro-C data in HFF cells.[4] To assign TAD lengths, each bead represented 1 kilobase pair (kbp) for simulations with $\overline{N_{TAD}} \approx 200$ beads. Beads in simulations with $\overline{N_{TAD}} \approx 100$ represented 2 kbp, and beads in simulations with $\overline{N_{TAD}} \approx 50$ represented 4 kbp. TADs were identified as in ref. (3) using the Arrowhead algorithm in Juicer tools v1.22.01[5] with 5 kbp resolution. Biological TAD lengths were placed in 20 logarithmically spaced bins, and simulated TAD lengths were chosen between 60 kbp and 800 kbp from this distribution (see Fig. S3(a) for an example). Anchors for the generated TAD lengths were placed on chains such that there were multiple TADs with potentially different lengths per chain, and each chain had different TAD lengths. We first generate a list of 3000 random TAD lengths from the distribution. Starting from the beginning of the list, we assign as many TADs as possible that can fit on a single chain with $N$ beads. Consider a melt with $N = 800$ beads per chain. If the first five TADs in the list have a cumulative length of 750 beads and the sixth TAD has a length of 80 beads, TADs one through five are placed on the first chain while the sixth TAD is assigned to the next chain. TADs are placed consecutively and centered along the chain contour (see Fig. S3(b)). This means that if the sum of TAD lengths is 750 beads on a chain with 800 beads, there are 25 beads on both chain ends that are not part of TADs. We continue this process until all chains have been assigned TADs (without exhausting the list of 3000). Figs. S3(c) and S3(d) show histograms for the number of TADs per chain and TAD coverage per chain (total number of beads within TADs) for an example of $\overline{N_{TAD}} \approx 200$ beads on chains with $N = 800$ beads. Table S2 shows combinations of $\overline{N_{TAD}}$ and $\lambda$ used in this work.

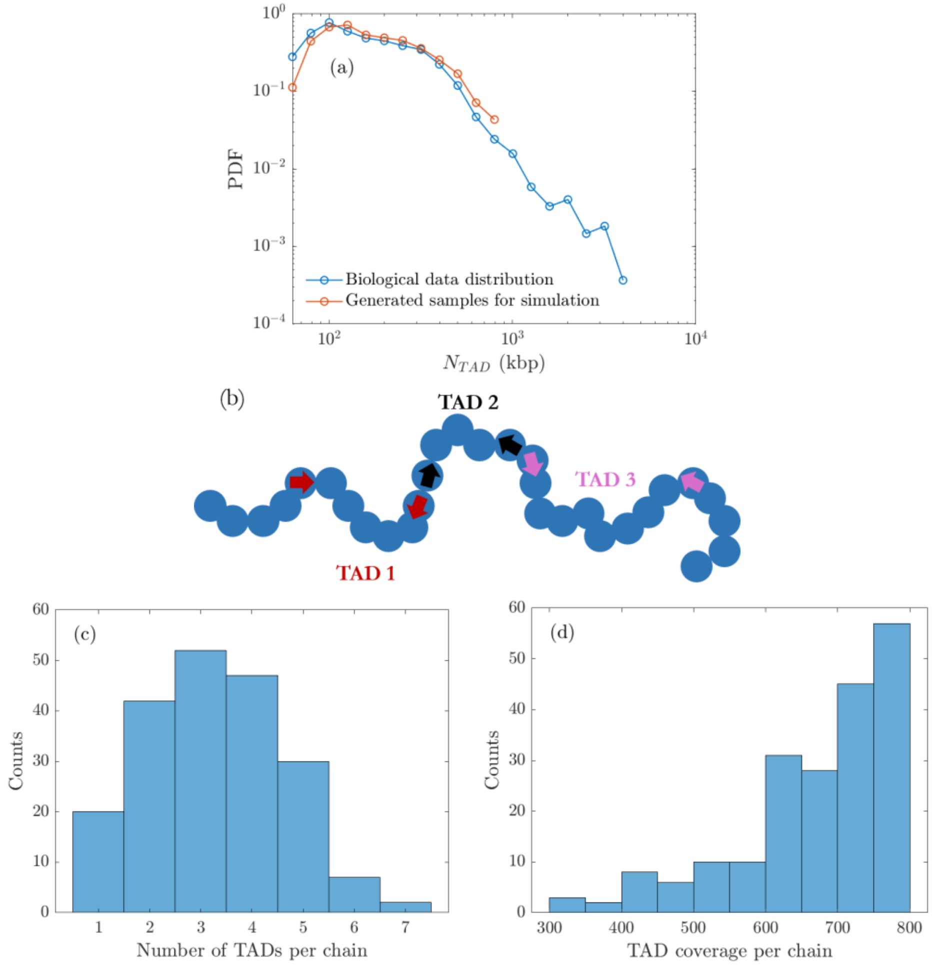


**Figure S3**: a) Distribution of TAD lengths from Micro-C data[4] (blue) compared to an example distribution of generated TAD lengths used for simulations (red). b) Schematic showing an example of TAD placement along a chain. Arrows represent TAD anchors. The anchors for each TAD are color coded (red, black, and pink). Left arrows only stop cohesin domains translocating from left to right; right arrows only stop cohesin translocating from right to left. TADs are centered along the chain contour such that there are four beads on both chain ends that are not within TADs. c) Example histogram of number of TADs per chain with $\overline{N_{TAD}} \approx 200$ beads and $N = 800$ beads per chain. d) Example histogram of TAD coverage per chain with $\overline{N_{TAD}} \approx 200$ beads and $N = 800$ beads per chain.

**Table S2**: Parameters for simulations with TADs

| $\overline{N_{TAD}}$ **(beads)** | **Uniform or distribution of TAD lengths** | $\lambda$ **(beads)** |
|---|---|---|
| 200 | Uniform | 200 |
| 200 | Distribution | 200 |
| 200 | Distribution | 50 |
| 100 | Uniform | 100 |
| 100 | Distribution | 100 |
| 50 | Uniform | 50 |
| 50 | Distribution | 50 |
| 50 | Distribution | 200 |

#### c. Ellipsoidal pervaded volumes and ellipsoidal radial distribution functions (eRDF)

Consider a chain segment of interest. We calculate the segment's center of mass (COM) and gyration tensor. The gyration tensor is characterized by eigenvalues $\lambda_1^2$, $\lambda_2^2$ and $\lambda_3^2$ with eigenvectors $\mathbf{v_1}$, $\mathbf{v_2}$, and $\mathbf{v_3}$ respectively. The coordinate of every bead in the melt is rewritten in the gyration tensor eigenbasis; that is, in a coordinate system with origin at the COM of the segment of interest, and using $\mathbf{v_1}$, $\mathbf{v_2}$, and $\mathbf{v_3}$ as the new basis vectors such that its coordinate is $(x_{eig,bead}, y_{eig,bead}, z_{eig,bead})$. We define the ellipsoidal pervaded volume as an ellipsoid with equation

$$\left(\frac{x_{eig}}{\lambda_1}\right)^2 + \left(\frac{y_{eig}}{\lambda_2}\right)^2 + \left(\frac{z_{eig}}{\lambda_3}\right)^2 = 1\,, \tag{S4}$$

where $x_{eig}$, $y_{eig}$, and $z_{eig}$ are written in the eigenbasis. Each bead can be described by

$$\left(\frac{x_{eig,bead}}{\lambda_1}\right)^2 + \left(\frac{y_{eig,bead}}{\lambda_2}\right)^2 + \left(\frac{z_{eig,bead}}{\lambda_3}\right)^2 = r_{e,bead}^2 \tag{S5}$$

such that the bead lies on the surface of an ellipsoid that is similar to the ellipsoidal pervaded volume of the segment of interest. We define the eRDF at a particular $r_e$ as the number of beads with $r_e \leq r_{e,bead} \leq r_e + \Delta r_e$ divided by the volume of an ellipsoidal shell with inner radii $r_e\lambda_1$, $r_e\lambda_2$, and $r_e\lambda_3$ and outer radii $(r_e + \Delta r_e)\lambda_1$, $(r_e + \Delta r_e)\lambda_2$, and $(r_e + \Delta r_e)\lambda_3$. The volume of this shell is $\frac{4\pi}{3}\lambda_1\lambda_2\lambda_3[(r_e + \Delta r_e)^3 - r_e^3]$.

#### d. Intra- and inter- TAD contact fractions

Intra- and inter- TAD contact fractions from Micro-C in two cell types/species, HFFs[4] and mESCs,[6] were calculated as described previously.[3] The intra-TAD contact fraction $F_{intra}(N_{TAD})$ is the fraction of all intra-chain contacts (or intra-chromosomal contacts for experimental data) made by a TAD with itself. The data shown in Fig. 5(b) of the main text are from active melt simulations with $N = 1600$ beads per chain, $\lambda \approx d \approx 200$ beads, and a distribution of TAD lengths with $\overline{N_{TAD}} \approx 200$ beads. Theoretical predictions for $F_{intra}(N_{TAD})$ and the inter-TAD contact fraction $F_{inter}(N_{TAD})$ were calculated by first considering the following function to describe contact probabilities:

$$P_{cross}(s) \approx s^{-\frac{3}{2}}\left[1 + \left(\frac{s}{g_{relax}}\right)^{\alpha_1}\right]^{\frac{3}{4\alpha_1}}\left[1 + \left(\frac{s}{A}\right)^{\alpha_2}\right]^{-\frac{3}{4\alpha_2}}\,, \tag{S6}$$

which has asymptotic scaling behavior

$$P_{asym}(s) \approx \begin{cases} s^{-\frac{3}{2}}\,, & s \leq g_{relax} \\ (s g_{relax})^{-\frac{3}{4}}\,, & g_{relax} < s \leq A \\ \left(\frac{A}{g_{relax}}\right)^{\frac{3}{4}} s^{-\frac{3}{2}}\,, & A < s \end{cases}. \tag{S7}$$

We fit the logarithm of experimental and simulation contact probability data to the logarithm of Eq. (S6) as in ref. (3) (see Table S3). The HFF and mESC data were fit simultaneously and normalized such that $P(5\text{kbp}) = 1$; the resulting parameters are the same as in ref. (3). Simulation data were fit for $1 \leq s \leq 1000$ beads. Figure S4 shows the simulation data compared to the fit.

**Table S3**: Parameters and 95% confidence intervals for fitting genome-wide or melt-wide contact probabilities to Eq. (S6).

| System | $\alpha_1$ | $\alpha_2$ | $g_{relax}$ | $A$ |
|---|---|---|---|---|
| Experiments | 1.7<br>(1.4, 2.1) | 2.7<br>(2.2 3.1) | 26 kbp<br>(25, 27) kbp | 350 kbp<br>(340, 370) kbp |
| Simulation | 2.5<br>(2.4, 2.7) | 3.9<br>(3.7, 4.0) | 15.2 beads<br>(15.1, 15.3) beads | 268 beads<br>(267, 270) beads |

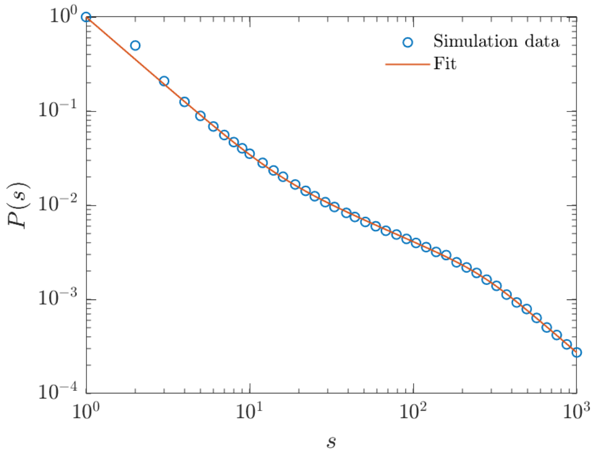


**Figure S4**: Melt-wide intra-chain contact probabilities for $N = 1600$ beads, $\lambda \approx d \approx 200$ beads, and a distribution of TAD lengths with $\overline{N_{TAD}} \approx 200$ beads. For clarity, 50 points uniformly spaced along the logarithmic axis are shown between $s = 1$ and $s = 1000$ beads. The red curve is the best fit to Eq. (S6) using parameters shown in Table S3.

The theoretical intra-TAD contact fraction $F_{intra}(N_{TAD})$ is estimated as

$$F_{intra}(N_{TAD}) \approx \frac{\int_1^{N_{TAD}} (N_{TAD} - s)\, P_{asym}(s) ds}{\int_1^{\frac{L}{2}} N_{TAD}\, P_{asym}(s) ds} \tag{S8}$$

where $P_{asym}(s)$ is given by Eq. (S7), $g_{relax}$ and $A$ are from Table S3, $N_{TAD}$ is the TAD length, and $L > A$ is the average chain (or chromosome) length. $L = 1600$ beads for simulations shown in Fig. 6(b) in the main text, and we take $L = 100$ megabase pairs for experimental data. The numerator is approximately the number of intra-TAD contacts, while the denominator is approximately the total number of contacts made by the TAD. Using Eq. (S7), the denominator, or the total number of contacts, is approximately

$$\begin{aligned} &\int_1^{\frac{L}{2}} N_{TAD}\, P_{asym}(s) ds \\ &= N_{TAD} \left[ \int_1^{g_{relax}} s^{-3/2}\, ds + \int_{g_{relax}}^{A} (s g_{relax})^{-3/4}\, ds + \int_A^{\frac{L}{2}} \left(\frac{A}{g_{relax}}\right)^{3/4} s^{-3/2} ds \right] \\ &= N_{TAD} \left[ 1 - \frac{1}{g_{relax}^{1/2}} + 2\left(\frac{A}{g_{relax}^3}\right)^{1/4} - \frac{2}{g_{relax}^{1/2}} + \left(\frac{A}{g_{relax}^3}\right)^{1/4} - \left(\frac{A}{g_{relax}}\right)^{3/4} \left(\frac{2}{L}\right)^{1/2} \right] \\ &\approx N_{TAD} \left[ 1 + \left(\frac{A}{g_{relax}}\right)^{3/4} \left(\frac{3}{A^{1/2}} - \left(\frac{2}{L}\right)^{1/2}\right) \right], \end{aligned} \tag{S9}$$

where in the last line we drop the smaller terms from the first two integrals. The numerator of Eq. (S8) has different forms depending on how the TAD length $N_{TAD}$ compares to $g_{relax}$ and $A$ (in all cases, $N_{TAD} > g_{relax}$). If $g_{relax} \leq N_{TAD} < A$, the last line of Eq. (S7) does not need to be considered and the numerator of Eq. (S8) is approximately

$$\begin{aligned} &\int_1^{N_{TAD}} (N_{TAD} - s)\, P_{asym}(s) ds \\ &= \int_1^{g_{relax}} (N_{TAD} - s) s^{-3/2}\, ds + \int_{g_{relax}}^{N_{TAD}} (N_{TAD} - s)(s g_{relax})^{-3/4}\, ds \\ &= N_{TAD} + \frac{8}{5} \frac{N_{TAD}^{5/4}}{g_{relax}^{3/4}} - \frac{3}{5} g_{relax}^{1/2} - 3 \frac{N_{TAD}}{g_{relax}^{1/2}} + 1 \\ &\approx N_{TAD} \left( 1 + \frac{8}{5} \frac{N_{TAD}^{1/4}}{g_{relax}^{3/4}} \right), \end{aligned} \tag{S10}$$

where the last line is approximate for $N_{TAD} \gg g_{relax} \gg 1$. If $g_{relax} < A \leq N_{TAD}$, the last line of Eq. (S7) does need to be considered. The numerator of Eq. (S8) is approximately

$$\begin{aligned} &\int_1^{N_{TAD}} (N_{TAD} - s)\, P_{asym}(s) ds \\ &= \int_1^{g_{relax}} (N_{TAD} - s)\, s^{-3/2} ds + \int_{g_{relax}}^{A} (N_{TAD} - s)(s g_{relax})^{-3/4}\, ds + \int_A^{N_{TAD}} (N_{TAD} - s) \left( \frac{A}{g_{relax}} \right)^{3/4} s^{-3/2} ds \\ &= N_{TAD} \left[ 1 + \left( \frac{A}{g_{relax}} \right)^{3/4} \left( \frac{3}{A^{1/2}} - \frac{1}{N_{TAD}^{1/2}} \right) - \frac{3}{g_{relax}^{1/2}} \right] - \left[ \left( \frac{A}{g_{relax}} \right)^{3/4} \left( 2N_{TAD}^{1/2} - \frac{3}{5} A^{1/2} \right) + \frac{3}{5} g_{relax}^{1/2} - 1 \right] \\ &\approx N_{TAD} \left( 1 + 3 \frac{A^{1/4}}{g_{relax}^{3/4}} \right) - \left( \frac{A}{g_{relax}} \right)^{3/4} \left( 2N_{TAD}^{1/2} - \frac{3}{5} A^{1/2} \right), \end{aligned} \tag{S11}$$

where the last line is approximate for $N_{TAD} \gg A \gg g_{relax} \gg 1$. To summarize, combining Eqs. (S8) - (S11), the theoretical intra-TAD contact fraction is approximately

$$F_{intra}(N_{TAD}) \approx \begin{cases} \dfrac{g_{relax}^{3/4} + \frac{8}{5} N_{TAD}^{1/4}}{g_{relax}^{3/4} + 3A^{1/4} - A^{3/4} \left( \frac{2}{L} \right)^{1/2}}, & g_{relax} \leq N_{TAD} < A \\ \dfrac{N_{TAD} \left[ g_{relax}^{3/4} + 3A^{1/4} \right] + \frac{3}{5} A^{5/4} - 2A^{3/4} N_{TAD}^{1/2}}{N_{TAD} \left[ g_{relax}^{3/4} + 3A^{1/4} - A^{3/4} \left( \frac{2}{L} \right)^{1/2} \right]}, & A \leq N_{TAD} \end{cases} \tag{S12}$$

and the inter-TAD contact fraction is $F_{inter}(N_{TAD}) \approx 1 - F_{intra}(N_{TAD})$.

**e. Apparent entanglement strand length**

Cohesin bonds were broken before quantifying apparent entanglement strands in simulated melts. The Z1+ package[7] was used to calculate the number of entanglements per chain $Z$. Since some chains in the active melts have no entanglements, we do not directly calculate the entanglement strand per chain as $N/Z$. Rather, we first calculate the average number of entanglements per chain throughout the melt $<Z>_{melt}$. The number of beads in an (apparent) entanglement strand is $N_e = N/<Z>_{melt}$, which is essentially the "modified S-kink" $N_e$ estimator in Z1+. Another reason why we use this estimator is that it does not rely on any assumption of the statistics of a chain's primitive path (meaning the primitive path does not necessarily need to be a random walk). The passive $N_e$ from our simulations of $\approx 36 \pm 1$ and $\approx$

$38 \pm 1$ for melts with 200 and 400 beads, respectively, are consistent with the literature value of S-kink $N_e$ between 35 and 39 for a monodisperse flexible linear melt with 100 chains, each with 400 beads.[7] Depending on the estimation method, the range of $N_e$ for monodisperse flexible linear melts is between 35 and 85.[7,8]

#### II. Extended data

##### a. Internal distances and contact probabilities

Figure S5 shows mean squared internal distances $<r^2(s)>$ and contact probabilities $P(s)$ for various active melts. Fig. S5(a) shows $<r^2(s)>$ for melts without TADs but with different $\lambda \approx d$, combining the data from Figs. 2(a) and 2(b) in the main text. In the compact regime, simulations without TADs and $\lambda \approx d \approx$100 or 200 (green and purple curves) are consistent with $<r^2(s)> \sim s^{2/7}$ and an apparent fractal dimension of $D \approx 7$. As discussed in the main text, we suggest that these melts are similar to fractal loopy globules (FLGs).[9] Simulations without TADs and $\lambda \approx d \approx$25 or 50 (blue and orange curves) appear to exhibit a scaling behavior closer to $<r^2(s)> \sim s^{1/2}$ in the compact regime, which would be consistent of $D \approx 4$. We suggest that this is because $\lambda \approx$25 and 50 are smaller than the true entanglement strand needed for FLG-like conformations to form. Note that the entanglement strand for passive flexible linear melts ranges between $N_e \approx 35$ and $N_e \approx 85$ depending on the estimation method.[7,8] Further clarification will be explored in future work, in addition to the dynamics of these systems. Fig. S5(c) displays the contact probabilities $P(s)$ for the same melts as in Fig. S5(a), which are all consistent with the mean-field estimate of $P(s) \sim s^{-3/D}$.

Figs. S5(b) and S5(d) compare $<r^2(s)>$ and $P(s)$, respectively, for active melts without TADs to active melts with a distribution of TADs with mean length either larger or smaller than $\lambda$. The conformations of melts with $\overline{N_{TAD}} > \lambda \approx d$ are similar to those without TADs (see orange squares compared to blue curves in Figs. S5(b) and S5(d)). This is because cohesins do not frequently extrude loops much longer than $\lambda$ so that these TADs have limited impact on chain conformations besides restricting the largest loops. On the other hand, when $\overline{N_{TAD}} < \lambda \approx d$, TAD anchors strongly suppress activity-induced compaction, as seen in the purple curves and circles compared to green curves in Figs. S5(b) and S5(d). In this case, TAD anchors restrict the size of most loops. These chains are still smaller than those in passive melts, reflecting the presence of loops smaller than $\overline{N_{TAD}}$.

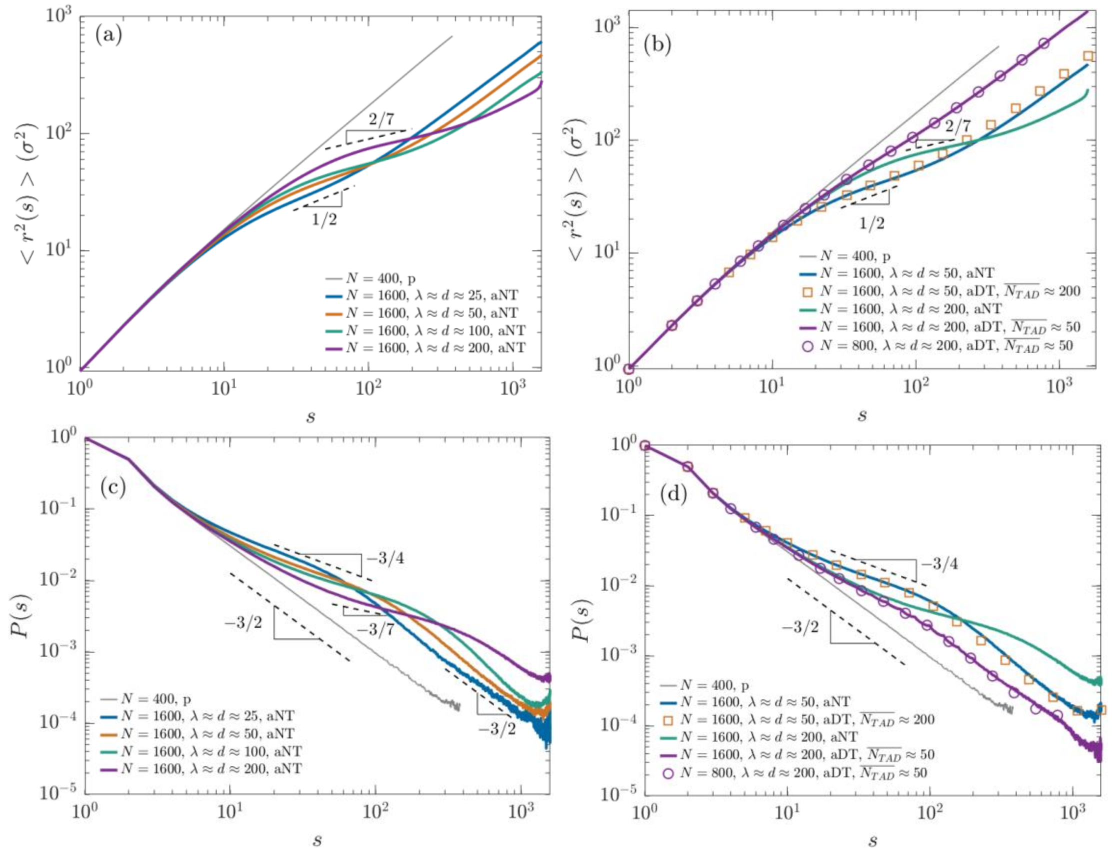


**Figure S5**: (a) Mean squared internal distances $<r^2(s)>$ for active melts without TADs. (b) Comparison of $<r^2(s)>$ for active melts without TADs to active melts with TADs with mean length either $\overline{N_{TAD}} > \lambda$ (blue curve compared to orange squares) or $\overline{N_{TAD}} < \lambda$ (green curve compared to purple curve and circles). (c) and (d) are the same as (a) and (b) but for contact probabilities $P(s)$. Dashed lines are guides to the eye representing various power laws.

#### b. Segment size distributions

Figure S6 shows normalized distributions of the sizes of chain segments with $s$ beads like in Figure 3(b) in the main text. Distributions shown here are zoomed in to $0 \leq x \leq 1.5$. For melts with $\lambda \approx d \approx \overline{N_{TAD}}$ in Fig. S6(a), the normalized distribution is slightly shifted to shorter distances for all $s > g_{relax}$ compared to passive melts. For melts without TADs in Fig. S6(b), the normalized distribution is also shifted to shorter distances for $g_{relax} < s < A$ (green curves). For longer segments with $s > A$, the short distances are slightly suppressed while the peak is slightly larger (purple curves). Recall that $A$ is the maximum segment length with the anomalously high apparent fractal dimension. The differences between active and passive melts are subtle; the overall shapes are similar. However, we reiterate that the average segment sizes are much smaller in active melts than passive melts for all cases in Fig. S6 (see Figs. 1 – 3 in the main text).

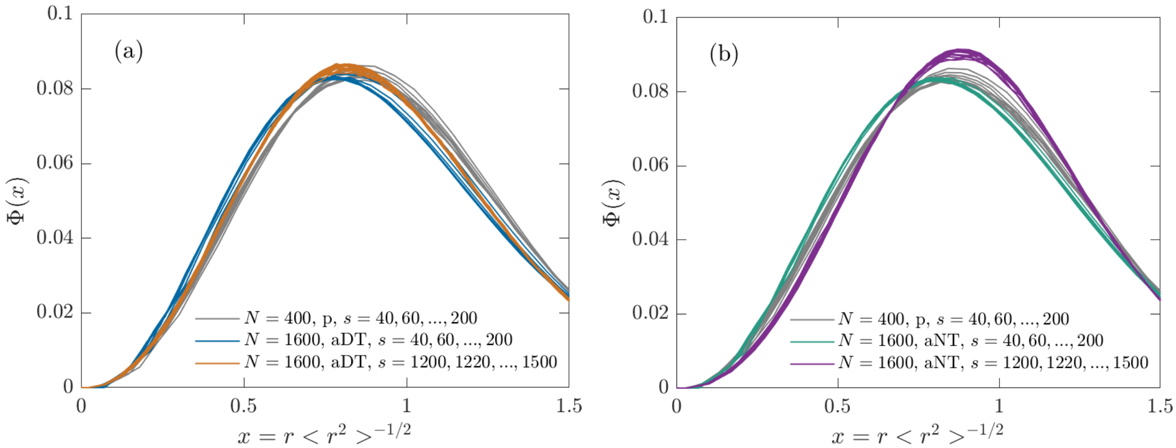


**Figure S6**: Normalized distributions of segments with varying number of beads in active melts with $\lambda \approx d \approx 200$, shown for $0 \leq x \leq 1.5$ where $x = r<r^2>^{-1/2}$. Full distributions are normalized such that $\int \Phi(x) 4\pi x^2 dx = 1$. (a) Comparing active melts with a distribution of TAD lengths to a passive melt. Blue curves are for $g_{relax} < s < A$ while orange curves are for $s > A$. (b) Comparing active melts without TADs to a passive melt. Green curves are for $g_{relax} < s < A$ while purple curves are for $s > A$.

#### c. Overlap parameters

Figure S7 shows the overlap parameter $O(s)$ for active melts with $\lambda \approx d \approx 100$ (Fig. S7(a)) and active melts with $\lambda \approx d \approx 50$ (Fig. S7(b)). For $\lambda \approx d \approx 100$, melts without TADs (blue curve) and with $\overline{N_{TAD}} \approx 100$ (orange curve) have non-monotonic $O(s)$ that first increase with $s$, then decrease, and increase once again. The decreasing regime of $O(s)$ without TADs is steeper than with TADs, with exponent between -4/7 and -3/4 which are the asymptotic limits for $D \approx 7$ and $D \approx 4$, respectively. With TADs of average length $\overline{N_{TAD}} \approx 100$, the decreasing regime is shallower than the expected asymptotic value of -3/4. This is because the overlap parameter increases for $s \geq 400$, so the range of the intermediate regime is likely not wide enough to reach the limiting scaling behavior. This is also potentially why the decreasing regime of $O(s)$ without TADs shows a power law slightly shallower than -4/7.

The shape of $O(s)$ for melts without TADs and $\lambda \approx d \approx 50$ (purple curve in Fig. S7(b)) is similar to the shape in melts without TADs and $\lambda \approx d \approx 100$ (blue curve in Fig. S7(a)). For $\lambda \approx d \approx 50$, the scaling behavior is consistent with $\sim s^{-1/4}$ in the compact regime where $O(s)$ is decreasing with $s$. This suggests a fractal dimension of $D \approx 4$ and is consistent with Fig. S5 above. Melts with $\overline{N_{TAD}} \approx 200 > \lambda$ have an $O(s)$ very similar to melts without TADs (green versus purple curve in Fig. S7(b)), again reflecting that large TADs only restrict the largest loops. Melts with $\overline{N_{TAD}} \approx \lambda \approx 50$ have monotonic $O(s)$ (orange curve in Fig. S7(b)), which is not the same behavior as melts with $\overline{N_{TAD}} \approx \lambda \approx 100$ and $\overline{N_{TAD}} \approx \lambda \approx 200$ in Fig. S7(a) and Fig. 4(b) in the main text, respectively. However, we suspect that the width of the intermediate regime is too narrow to see the limiting scaling behaviors.

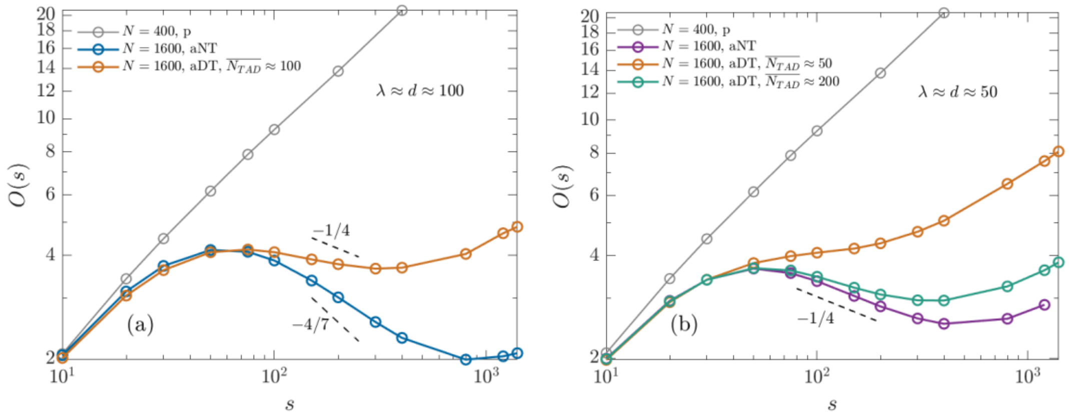


**Figure S7**: Overlap parameter for various active melts with and without TADs. Dashed lines are guides to the eye. (a) Melts with $\lambda \approx d \approx 100$ beads. (b) Melts with $\lambda \approx d \approx 50$ beads.

Figure S8 shows the overlap parameter for various active melts with (open squares) and without TADs (open circles) that have non-monotonic $O(s)$. The abscissa in Fig. S8(a) is normalized by $g_{relax}$, showing that the local maximum of $O(s)$ is at approximately $5g_{relax}$ for all cases. The abscissa in Fig. S8(b) is normalized by either $4\overline{N_{TAD}}$ if $\overline{N_{TAD}} \approx \lambda$ or $8\lambda$ if $\overline{N_{TAD}} > \lambda$ or there are no TADs. This normalization suggests the local minimum of $O(s)$ occurs at approximately $4\overline{N_{TAD}}$ or $8\lambda$ depending on the presence of TADs and their average length.

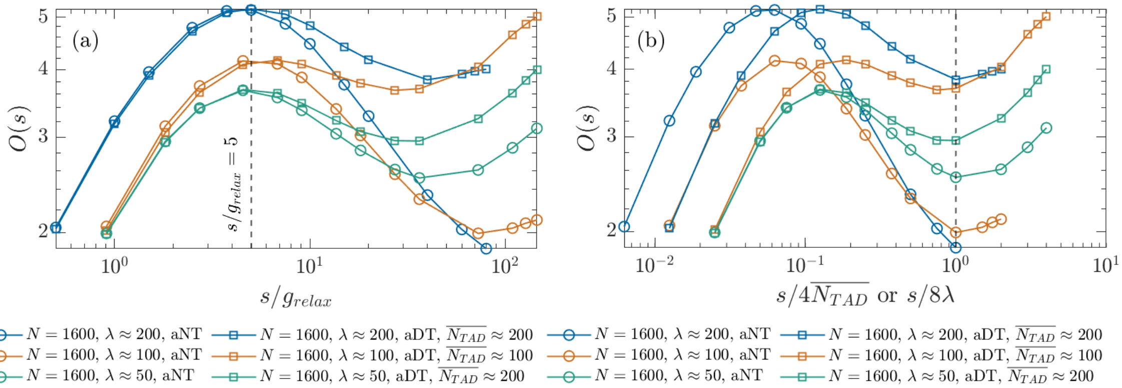


**Figure S8**: (a) Overlap parameter $O(s)$ with abscissa normalized by $g_{relax}$. (b) Overlap parameter $O(s)$ with abscissa normalized by either $4\overline{N_{TAD}}$ or $8\lambda$. All melts without TADs (open circles) were normalized by $8\lambda$. Two melts with TADs were normalized by $4\overline{N_{TAD}}$ (blue and orange open squares), while one was normalized by $8\lambda$ (green open squares).

##### d. Anisotropy, $r_{e,1/2}$, and ellipsoidal overlap parameter

Figure S9(a) shows the anisotropy of TADs from simulations with $N = 1600$ beads per chain, $\lambda \approx d \approx 200$, and a distribution of TAD lengths with $\overline{N_{TAD}} \approx 200$ (related to Fig. 5 in the main text). Anisotropy is defined as

$$Anisotropy = \frac{1}{2}\left[3\frac{Tr(\mathbf{S}^2)}{Tr(\mathbf{S})^2} - 1\right], \quad \text{(S13)}$$

where $S$ is the gyration tensor of the TAD of interest and $Tr$ signifies the matrix trace. Note that in some sources, this descriptor is called "asphericity". Anisotropy is 0 for a spherically symmetric object. Three-dimensional random walks like a Gaussian linear chain have anisotropy of approximately 0.53 (the limiting value for infinite-dimensional random walks is 0.4).[10] TAD anisotropy decreases with TAD length from ≈0.4 to ≈0.1 and is smaller than the value for Gaussian linear chains but still nonzero. The ratio $\lambda_1/\lambda_3$ decreases from ≈3.1 to ≈1.8, where $\lambda_1$ and $\lambda_3$ are the square roots of the largest and smallest gyration tensor eigenvalues, respectively. Figure S9(b) shows $r_{e,1/2}$ for TADs longer than 200 beads. This value denotes the normalized ellipsoidal "radii" (section I.c. above) where the "self" and "other" components of the ellipsoidal radial distribution functions become equal ($eRDF_s$=$eRDF_o$=0.5). That is, the number density of beads in ellipsoidal shells with inner radii smaller than $r_{e,1/2}\lambda_1$, $r_{e,1/2}\lambda_2$, and $r_{e,1/2}\lambda_3$ is dominated by beads from a TAD itself, while the density in shells with larger radii is dominated by other beads. $\lambda_1$, $\lambda_2$, and $\lambda_3$ are the square roots of the TAD's gyration tensor eigenvalues. Figure S9(c) shows the overlap parameter $O(s)$ calculated by considering either spherical pervaded volumes or ellipsoidal pervaded volumes (see discussion of Fig. 3 in main text and section I.c. above). The shapes of both methods are similar, though the calculation using ellipsoidal volumes results in fewer overlaps.

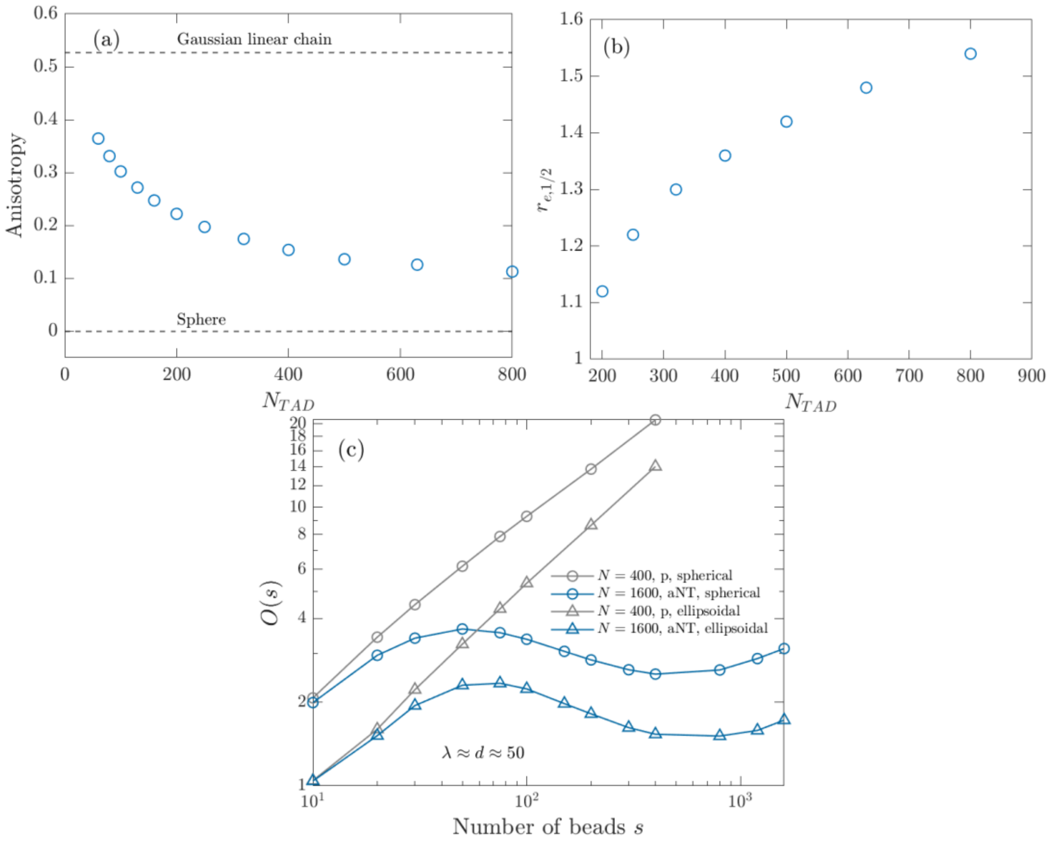


**Figure S9**: (a) Anisotropy of TADs in simulations with $N = 1600$ beads per chain, $\lambda \approx d \approx 200$, and a distribution of TAD lengths with $\overline{N_{TAD}} \approx 200$. (b) Normalized ellipsoidal radii where $eRDF_s$=$eRDF_o$=0.5. (c) Comparison of overlap parameters calculated using spherical and ellipsoidal pervaded volumes.

#### e. Apparent entanglement strand

Figure S10(a) shows the apparent $N_e$ for active melts with $\lambda \approx d \approx 50$ beads with and without TADs. Without TADs (orange circles), the apparent $N_e$ is almost linear with chain length, just like for the active melts without TADs and $\lambda \approx d \approx 100$ or $\lambda \approx d \approx 200$ as shown in Fig. 7 in the main text. Simulations with a distribution of TAD lengths or uniform TAD lengths with $\overline{N_{TAD}} \approx 50$ (orange triangles and plus signs, respectively) have smaller $N_e$ than without TADs but still larger than the passive case. Melts with a distribution of TAD lengths with $\overline{N_{TAD}} \approx 200 > \lambda$ (orange crosses) have apparent $N_e$ in between the cases without TADs and $\overline{N_{TAD}} \approx 50 \approx \lambda$. This is because TAD anchors mainly function to limit loop locations, and the effect is not as pronounced when TADs are longer than $\lambda$. Figure S10(b) shows the ratio $N/N_e$ for melts without TADs, showing that there are fewer than 3.5 entanglements per chain in all cases studied. Figure S10(c) shows linear fits to the apparent $N_e$ as a function of $N$ for the same melts without TADs.

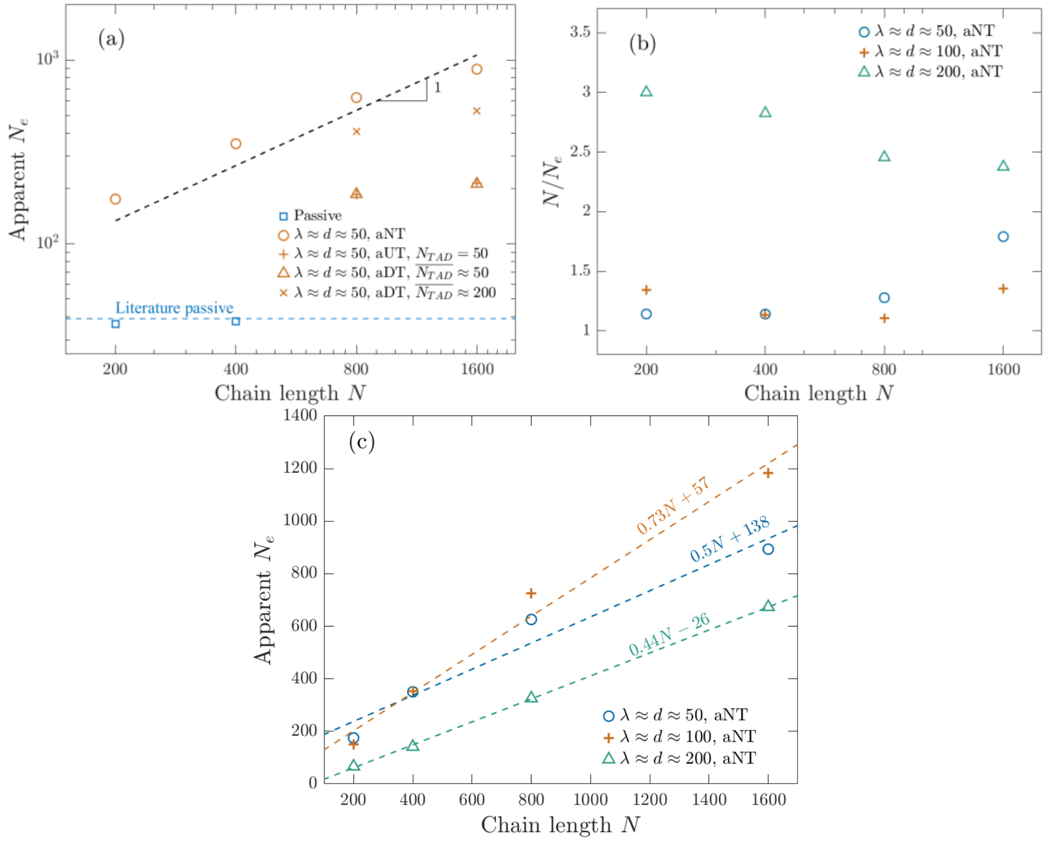


**Figure S10**: (a) Apparent $N_e$ for passive melts and active melts with $\lambda \approx d \approx 50$ beads with and without TADs. The dashed line is a guide to the eye for $N_e \sim N^1$. (b) Number of entanglements per strand for active melts without TADs. (c) Linear fits of apparent $N_e$ as a function of chain length $N$ for active melts without TADs.
